# Intranasal administration of a monoclonal neutralizing antibody protects mice against SARS-CoV-2 infection

**DOI:** 10.1101/2021.06.09.447662

**Authors:** Sandro Halwe, Alexandra Kupke, Kanika Vanshylla, Falk Liberta, Henning Gruell, Matthias Zehner, Cornelius Rohde, Verena Krähling, Michelle Gellhorn-Serra, Christoph Kreer, Michael Klüver, Lucie Sauerhering, Jörg Schmidt, Zheng Cai, Fei Han, David Young, Guangwei Yang, Marek Widera, Manuel Koch, Anke Werner, Lennart Kämper, Nico Becker, Michael S Marlow, Markus Eickmann, Sandra Ciesek, Felix Schiele, Florian Klein, Stephan Becker

**Affiliations:** Institute of Virology, Philipps University Marburg, Hans-Meerwein-Straße 2, 35043 Marburg, Germany; German Center for Infection Research (DZIF), partner site Giessen-Marburg-Langen, Marburg, Germany; Institute of Virology, Faculty of Medicine and University Hospital Cologne, University of Cologne, 50931 Cologne, Germany; German Center for Infection Research (DZIF), partner Site Bonn-Cologne, 50931 Cologne, Germany; Center for Molecular Medicine Cologne (CMMC), University of Cologne, 50931 Cologne, Germany; Biotherapeutics Discovery, Boehringer Ingelheim Pharma GmbH & Co. KG, Birkendorfer Strasse 65, 88397 Biberach an der Riss, Germany; Biotherapeutics Molecule Discovery, Boehringer Ingelheim Pharmaceuticals Inc., Ridgefield, CT, USA; Institute for Medical Virology, University Hospital Frankfurt, Goethe University Frankfurt am Main, 60596, Germany; German Center for Infection Research (DZIF), partner site Frankfurt am Main, Germany; Fraunhofer Institute for Molecular Biology and Applied Ecology (IME), Branch Translational Medicine and Pharmacology, Frankfurt am Main, 60596, Germany; Institute for Dental Research and Oral Musculoskeletal Biology and Center for Biochemistry, University of Cologne, 50931 Cologne, Germany

**Author notes:** Corresponding author: Stephan Becker, Institut für Virologie, Philipps-Universität Marburg, Hans-Meerwein-Straße 2, 35043 Marburg, Germany, Tel.: +49 6421/28-66254, Fax.: +49 6421/28-68962. **Author Contributions** Conceptualization: SH, AK, SB Methodology: SH, AK, CR, VK, SB, MZ, HG, KV, CK, FL, ZC, FH, DY, GY, MSM, FS Investigation: SH, CR, VK, LS, MGS, MK, JS, AW, ME, NB, LK, MZ, HG, KV, CK, MK, FL, ZC, FH, GY, MSM, FS Visualization: SH, MZ, HG, KV, FL, ZC, GY Funding acquisition: SB, FK Project administration: SB, FK, ME, MSM, FS Supervision: SB, AK, FK, CK, MSM, FS Writing – original draft: SH, SB, KV, MZ, FL, ZC, FH, GY Writing – review & editing: all authors. **Competing interests** A patent application encompassing DZIF-10c has been filed by the University of Cologne, listing F.K., S.B., C.K., M.Z., and H.G. as inventors (20182325.9).

**Keywords:** SARS-CoV-2, monoclonal antibody, neutralizing antibody, virus, animal experiments, mice, transduction, intranasal administration, topical administration

## Abstract

Despite recent availability of vaccines against severe acute respiratory syndrome coronavirus type 2 (SARS-CoV-2), there is an urgent need for specific anti-SARS-CoV-2 drugs. Monoclonal neutralizing antibodies are an important drug class in the global fight against the SARS-CoV-2 pandemic due to their ability to convey immediate protection and their potential to be used as both, prophylactic and therapeutic drugs. Clinically used neutralizing antibodies against respiratory viruses are currently injected intravenously, which can lead to suboptimal pulmonary bioavailability and thus to a lower effectiveness.

Here we describe DZIF-10c, a fully human monoclonal neutralizing antibody that binds the receptor-binding domain of SARS-CoV-2 spike protein. DZIF-10c displays an exceptionally high neutralizing potency against SARS-CoV-2 and retains activity against the variants of concern B.1.1.7 and B.1.351. Importantly, not only systemic but also intranasal application of DZIF-10c abolished presence of infectious particles in the lungs of SARS-CoV-2 infected mice and mitigated lung pathology. Along with a favorable pharmacokinetic profile, these results highlight DZIF-10c as a novel human SARS-CoV-2 neutralizing antibody with high *in vitro* and *in vivo* antiviral potency. The successful intranasal application of DZIF-10c paves the way for clinical trials investigating topical delivery of anti-SARS-CoV-2 antibodies.

**Significance Statement:** Monoclonal neutralizing antibodies are important in the global fight against the SARS-CoV-2 pandemic due to their ability to convey immediate protection. However, their intravenous application might lead to suboptimal bioavailability in the lung. We here precisely characterize a new monoclonal neutralizing antibody (DZIF-10c) that binds to the receptor binding domain of the spike protein of SARS-CoV-2. DZIF-10c neutralizes SARS-CoV-2 with exceptionally high potency and maintains activity against circulating variants of concern. The antibody has a favorable pharmacokinetic profile and protects mice from SARS-CoV-2 infection. Importantly, we show that intranasal administration of DZIF-10c generates protective efficacy. These results not only identify DZIF-10c as a novel highly potent neutralizing antibody, but further pave the way for a topical application of anti-SARS-CoV-2 antibodies.

## Main Text

### Introduction

The pandemic spread of severe acute respiratory syndrome coronavirus type 2 (SARS-CoV-2) poses unprecedented challenges for global public health systems. Facing more than 164 Mio. Coronavirus disease 2019 (COVID-19) cases and 3.4 Mio. fatalities until 19^th^ May 2021 (1), governments worldwide have enacted massive non-pharmaceutical countermeasures to mitigate the pandemic with drastic side effects for the global economy and daily live. Although the successful development of several vaccines will reduce the number of new SARS-CoV-2 infections (2–4), there is still an urgent need for antiviral interventions to prevent and treat COVID-19. In recent years, monoclonal neutralizing antibodies (nAbs) targeting viral surface proteins have been proven to be effective antiviral interventions against viruses such as respiratory syncytial virus (RSV), Zaire ebolavirus or human immunodeficiency virus 1 (HIV-1) (5– 9). The spike (S) protein of SARS-CoV-2 has essential functions within the viral replication cycle mediating binding to the cellular receptor human angiotensin-converting enzyme 2 (hACE2), as well as fusion with the target cell’s endosomal membrane prior to nucleocapsid release into the cytoplasm. A majority of nABs against SARS-CoV-2 exert their antiviral properties by disrupting the interaction of S with its receptor hACE2 and thereby preventing viral entry (10, 11).

Several S-specific nAbs have been described to efficiently neutralize SARS-CoV-2 *in vitro* and *in vivo* (12– 19). The efficacy of two S-specific nAbs were already successfully tested in clinical phase III trials and gained emergency use authorization by the US government for the treatment of ambulatory patients with mild to moderate COVID-19 (14, 15). Despite these promising results, all S-specific nAbs available so far have to be administered via intravenous infusion, which is one reason why nAb therapy is cost-intensive and challenging in terms of patient management and compliance. Furthermore, the systemic application of an antibody might be suboptimal with regard to its bioavailability in the lung, the primary site-of-action against respiratory viruses such as SARS-CoV-2 (20). In order to make nAb therapy more feasible for COVID-19 treatment, it is of great interest not only to identify new potent nAbs but also to investigate alternative approaches for their administration.

Previously, we described the isolation of a large panel of monoclonal SARS-CoV-2 nAbs from twelve SARS-CoV-2-convalescent individuals (21). One of these nAbs (HbnC3t1p1_F4) showed high neutralizing capacity along with a favorable biochemical profile for large scale production and clinical use. For the current study, the C-terminal heavy chain lysine of HbnC3t1p1_F4 was removed to reduce potential charge heterogeneity (22), resulting in a slightly modified antibody named *DZIF-10c*. We show here by ELISA and surface plasmon resonance (SPR) that DZIF-10c binds the receptor-binding-domain (RBD) of SARS-CoV-2 S with nanomolar affinity. DZIF-10c efficiently neutralizes both, SARS-CoV-2 pseudoviruses as well as authentic SARS-CoV-2 with 100% inhibitory concentrations (IC_100_) of 0.01 µg/ml. Furthermore, prophylactic application of DZIF-10c *in vivo* efficiently prevents SARS-CoV-2 infection and markedly mitigates lung pathology in hACE2-transduced mice after both, systemic (intraperitoneal, i.p.) and topical (intranasal, i.n.) administration. These results highlight DZIF-10c as a very potent nAb with favorable *in vitro* characteristics as well as high *in vivo* efficacy against SARS-CoV-2 after i.p. and i.n. administration.

## Results

### DZIF-10c displays an extraordinary neutralizing capacity against SARS-CoV-2 and remains active against SARS-CoV-2 variants B.1.1.7 and B.1.351

To compare the characteristics of DZIF-10c with other antibodies that have already shown clinical efficacy in phase 3 studies (14), we analyzed the binding of DZIF-10c and REGN10933 as well as REGN10987 to the RBD of SARS-CoV-2 S (Fig. 1A). All three antibodies showed specific binding to the RBD of SARS-CoV-2 S by ELISA with half-maximal effective concentrations (EC_50_) between 0.046 µg/ml (DZIF-10), 0.057 µg/ml (REGN10933) and 0.061 µg/ml (REGN10987). S-binding was further confirmed by ELISA against the full trimeric S ectodomain, a truncated N-terminal S1 subunit as well as a monomeric S ectodomain while no binding to the unrelated Zaire ebolavirus glycoprotein was observed (Fig. S1). Using SPR we could further show that DZIF-10c targets the RBD of SARS-CoV-2 S with high affinity indicated by an equilibrium dissociation constant (K_D_) of 1.09 ± 0.22 nM (Fig. S2). As these results highlight DZIF-10c as a very potent S-binding antibody, we further characterized its neutralizing properties. The *in vitro* neutralizing activity of DZIF-10c was first evaluated against a panel of six pseudovirus variants (23, 24). DZIF-10c demonstrated potent neutralizing activity against all SARS-CoV-2 S variants tested, including a variant carrying the D614G mutation, with an average IC_50_ of 0.007 µg/ml (Fig. 1B). In line with these results, DZIF-10c efficiently neutralized authentic SARS-CoV-2 (BavPat1, lineage B.1) with a mean IC_100_ of 0.01 µg/ml (Fig. 1C, D). Since the emergence of several SARS-CoV-2 variants of concern (VOC) showed that certain amino acid changes (e.g. E484K, N501Y) in the RBD of the S protein may completely abolish neutralization of antibodies raised again the Wuhan strain of SARS-CoV-2 (25–29), we further tested the ability of DZIF-10c to neutralize pseudovirus particles bearing S proteins with single point mutations in the Wuhan background, the 69-70 deletion mutant, as well as authentic VOCs. The pseudovirus neutralization assay revealed that the neutralization capacity of DZIF-10c was not affected by 16 out of 19 tested point mutations in the RBD. Although DZIF-10c was able to efficiently neutralize pseudovirus particles bearing the B.1.1.7 spike variant, reduced neutralizing activity was observed against the E484K and F490S mutants and the B.1.351 variant as demonstrated by an about 2.5-log-fold reduction in the IC_50_ (Fig. S3, Fig. 1D). Importantly, activity of DZIF-10c was unchanged against the currently circulating authentic VOC B.1.1.7 and, though reduced by 17-fold, also active against VOC B.1.351 (Fig. 1C, D).

**Fig. 1:**
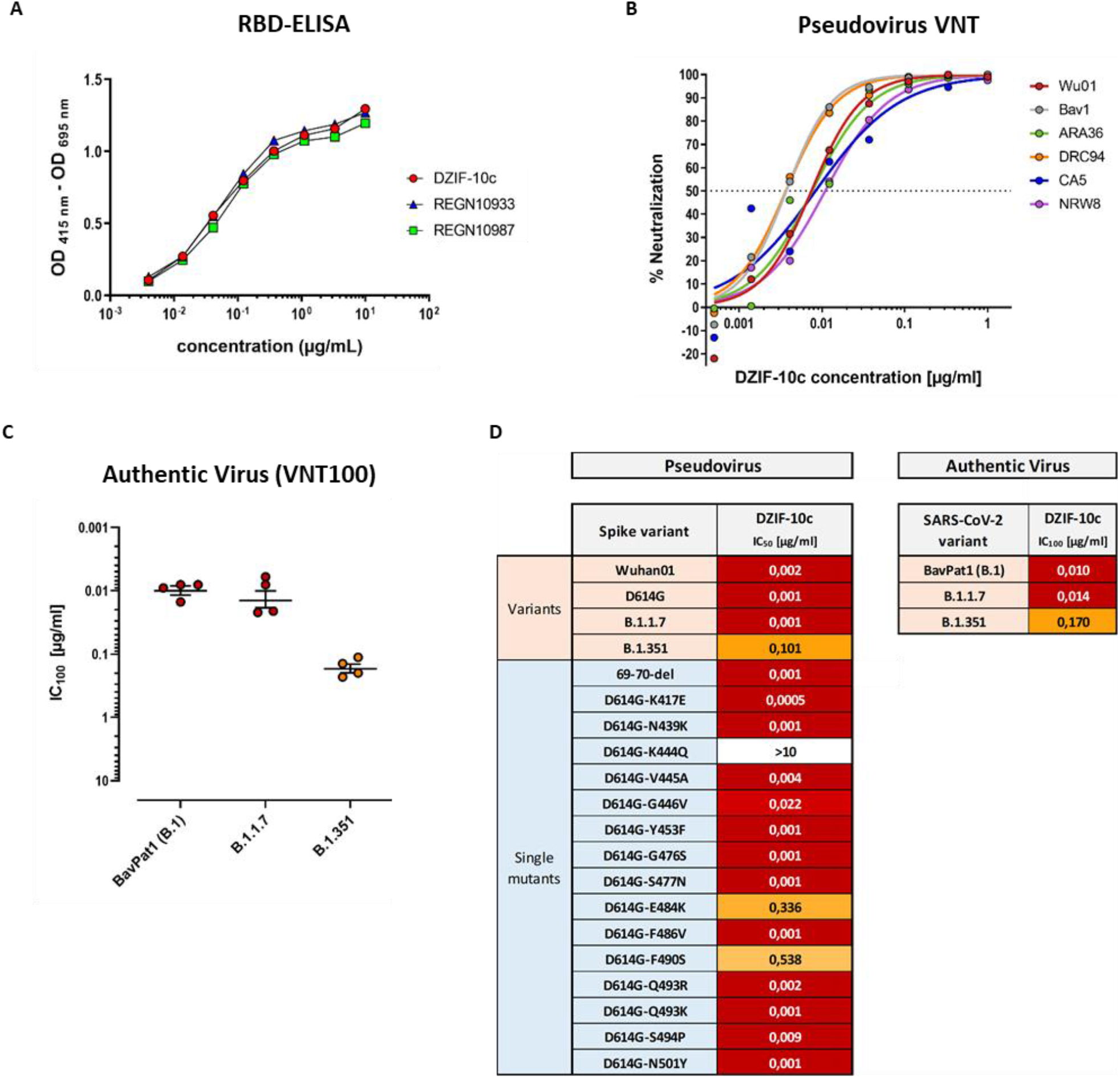
DZIF-10c binds and neutralizes SARS-CoV-2 with high potency. **(A)** Interaction of DZIF-10c, REGN10933 and REGN10987 with SARS-CoV-2 S RBD measured by ELISA. **(B)** Neutralizing activity of DZIF-10c against SARS-CoV-2 pseudoviruses bearing S proteins from different circulating strains. The dotted line indicates 50% neutralization (IC_50_). **(C)** Neutralization of authentic SARS-CoV-2 (BavPat1, B.1) and SARS-CoV-2 variants B.1.1.7 and B.1.351 by DZIF-10c measured by VNT100. Neutralization was defined as complete inhibition of CPE (IC_100_). Circles represent geometric means of four independent experiments. Lines and error bars indicate the overall mean with SEM. **(D)** Summary of the neutralizing activity (IC_50_ or IC_100_) of DZIF-10c against SARS-CoV-2 pseudoviruses bearing various mutations in the S protein as well as against authentic SARS-CoV-2.

### Structural analysis indicates binding of DZIF-10c to the prefusion conformation of S adjacent to the receptor binding motif

To investigate the structural basis of the neutralizing activity of DZIF-10c, the protein structure of the DZIF-10c Fab/SARS-CoV-2 S complex was determined using cryo-electron microscopy (cryo-EM) at a global resolution of 3.7 Å according to the 0.143 criteria(30) (Fig. S4B, C). The reconstructed 3D cryo-EM map of the analyzed Fab-antigen complex clearly shows the trimeric shape of the S protein and additional density for a single DZIF-10c Fab, bound to one of the three S protein protomers (Fig. 2A). The reconstructed S protein is arranged in a conformation showing one RBD in “up”-conformation, while the remaining two RBDs reside in “down”-conformation corresponding to the prefusion conformation (31). The DZIF-10c Fab fragment binds to the RBD in “up”-conformation and is arranged nearly perpendicular, relative to the symmetry axis of the S protein trimer (Fig. 2A). No Fab density was observed close to the remaining two RBDs in “down”-conformation, despite the 3:1 molar excess of Fab to S protein trimer suggesting that DZIF-10c preferably binds the “up”-conformation of the RBD, which corresponds to the activated prefusion state of the S protein. Furthermore, the data show that the approximated binding position and angle of DZIF-10c do not or only peripherally interfere with the ACE2 binding motif on the RBD (Fig. 2B), pointing to a mode of inhibition independent from directly blocking the ACE2 binding.

**Fig. 2:**
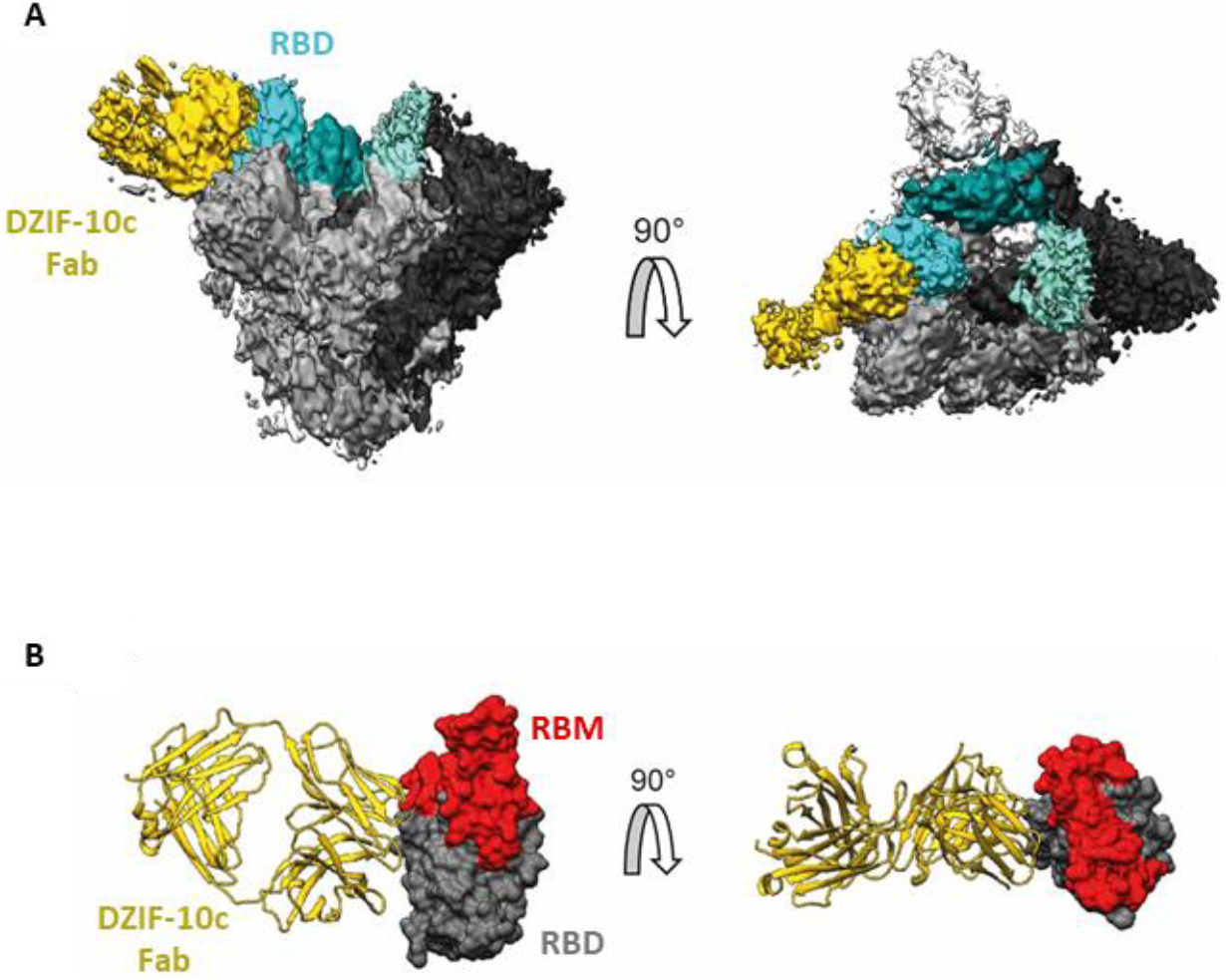
Cryo-EM map of the DZIF-10c Fab/SARS-CoV-2 S complex. **(A)** Cryo-EM map with S protein colored in gray shades (according to three protomers), density corresponding to the RBDs colored in cyan shades and density corresponding to the Fab fragment colored in yellow. left: side view; right: top view. **(B)** Approximate binding position and angle of DZIF-10c relative to the RBD. DZIF-10c colored in yellow, RBD (PDB-6XDG) colored in grey and receptor binding motif (RBM) on RBD colored in red.

### DZIF-10c shows a favorable pharmacokinetic profile *in vivo*

In view of its promising *in vitro* properties, we next characterized the pharmacokinetic profile of DZIF-10c after a single i.v. injection in two different mouse models. First, DZIF-10c was investigated in severe combined immunodeficient (SCID) mice expressing a human alpha chain Fc molecule instead of the endogenous mouse Fcgrt (Fig. 3A). The human neonatal Fc receptor (huFcRn) reduces lysosomal degradation of human IgG and plays a key role in antibody half-life. Mice genetically engineered to express the human neonatal Fc receptor are used as a surrogate model for antibody pharmacokinetics in humans (32). After administration of an antibody dose of 0.5 mg in PBS, antibody serum levels were determined by a human IgG ELISA using purified human myeloma IgG as on-plate standard. In these mice, DZIF-10c showed a favorable pharmacokinetic profile similar to two HIV-1-neutralizing human IgG antibodies that are currently in clinical investigation and show *in vivo* half-life of approximately 2 weeks (3BNC117) to 3 weeks (10-1074) in humans(7, 33, 34). In addition, DZIF-10c was administered to immunodeficient NRG mice that do not express the IL-2 receptor common gamma chain, carry a knock-out mutation in the *Rag1* gene, and do not develop murine lymphocytes or NK cells (Fig. 3B). This model has previously been used to faithfully reproduce the overall pharmacokinetic characteristics of antiviral antibodies in humans (6, 7, 34, 35). Again, compared to the reference antibodies the pharmacokinetic profile of DZIF-10c was similar or prolonged. Neither mouse model showed accelerated clearance and/or serum elimination of DZIF-10c.

**Fig. 3:**
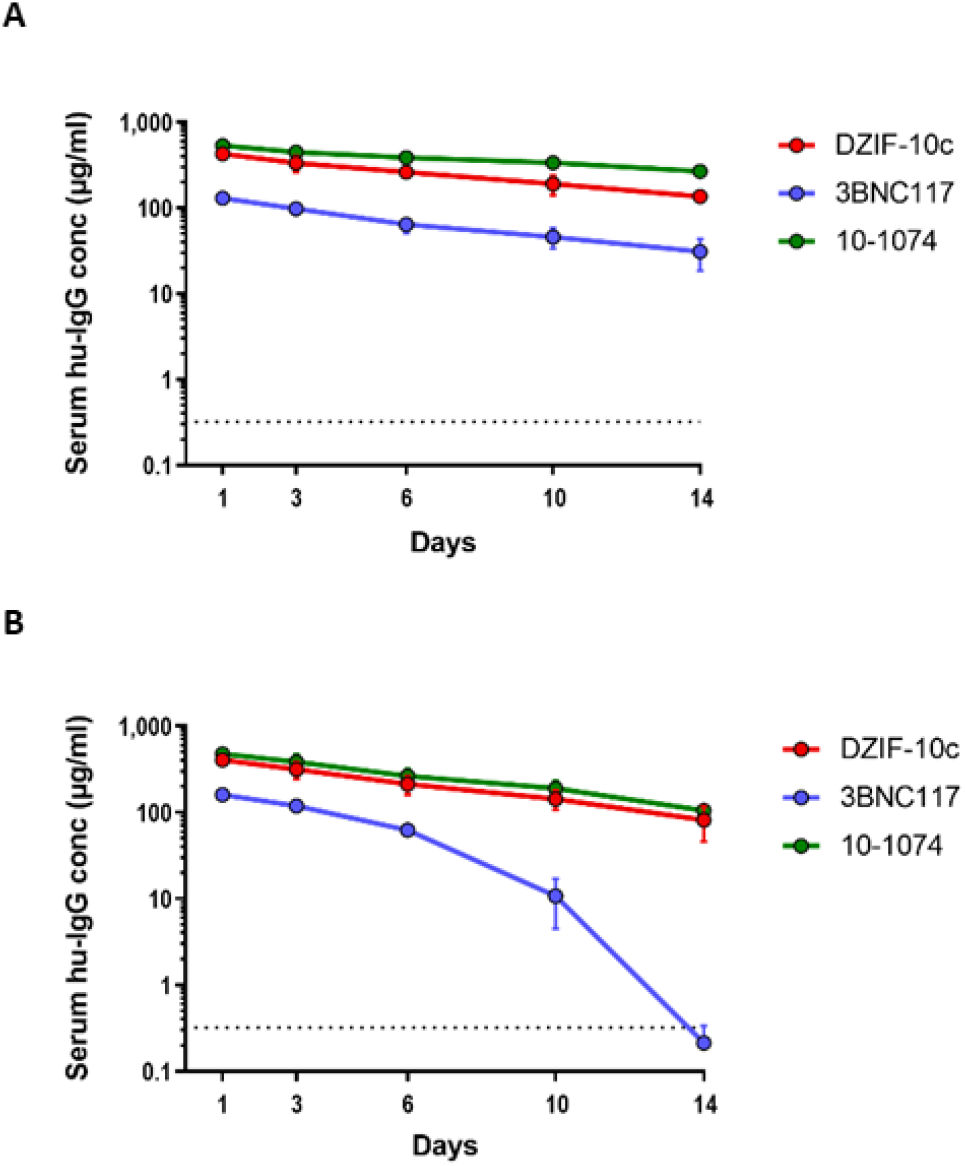
DZIF-10c shows a favorable pharmacokinetic profile in huFcRn and NRG mice. **(A)** Pharmacokinetic profile of DZIF-10c and two human anti-HIV-1 IgG1 antibodies in mice expressing the human neonatal Fc receptor after i.v. injection of a 0.5 mg dose. **(B)** Pharmacokinetic profile of DZIF-10c and two human anti-HIV-1 IgG1 antibodies in NRG mice after i.v. injection of a 0.5 mg dose. Antibody levels were determined by human IgG-specific ELISA.

### DZIF-10c efficiently protects hACE2-transduced mice from infection with SARS-CoV-2

To demonstrate the antiviral efficacy *in vivo*, we analyzed the prophylactic and therapeutic potential of DZIF-10c after SARS-CoV-2 challenge in a hACE2 transduction mouse model. Since mice are naturally not susceptible to SARS-CoV-2 infection due to incompatibility of the S protein to murine ACE2 (36–38), we transduced BALB/c mice intratracheally with an adenoviral vector carrying the genetic information for hACE2 and the reporter mCherry (Ad_ACE2-mCherry). This approach is based on a previously published model for middle east respiratory syndrome coronavirus (MERS-CoV) and has been shown to be suitable for modeling SARS-CoV-2 infection (39, 40) as well as preclinical testing of a SARS-CoV-2 vaccine candidate (41). Moreover, this model was used successfully by several other groups to study SARS-CoV-2 pathogenesis and antiviral interventions(18, 42, 43). The recombinant adenovirus additionally encodes the fluorescent protein mCherry, which allows the assessment of the transduction efficiency (Fig. S6). First, we assessed if DZIF-10c can protect hACE2-transduced mice from SARS-CoV-2 infection in a prophylactic setting. To this end, we transduced BALB/c mice with Ad_ACE2-mCherry on day three prior to challenge followed by a single dose of 40 mg/KG body weight DZIF-10c or an IgG isotype control antibody on day one prior to challenge (Fig. 4A). The antibodies were administered by two different routes to model a systemic (i.p.) or a topical (i.n.) administration. One day after treatment, mice were challenged with SARS-CoV-2 and monitored daily for changes in body weight, behavior and appearance. Four days post challenge, the mice were euthanized, and the lungs were collected for viral load determination as well as histological analyses. Neither significant body weight changes nor clinical symptoms were observed in any of the groups (Fig. S7), which is in line with previous reports of hACE2-transduced mice (41, 44). Importantly, this result indicated that the presence of DZIF-10c did not induce antibody-dependent enhancement (ADE). Regarding viral load, we observed high titers of infectious SARS-CoV-2 in lung homogenates of all mice that received the IgG isotype control antibody with 2.46×10^3^ (± 1.06×10^3^) TCID_50_/25 mg tissue in the i.n. group and 8.35×10^3^ (± 3.76×10^3^) TCID_50_/25 mg tissue in the i.p. group (Fig. 4B). In contrast, no infectious virus was detected in mice prophylactically treated with DZIF-10c independent on the route of delivery, indicating efficient neutralization of SARS-CoV-2 *in vivo*. In line with these results, viral genomic RNA (gRNA) measured by RT-qPCR was reduced significantly by approximately two log scales in DZIF-10c-treated mice (Fig. 4C). Intriguingly, in animals receiving the antibody i.n., the decrease was approximately three times greater compared to animals of the i.p. group. Because of the possibility that gRNA measured by RT-qPCR is partially derived either from input virus or intact, but neutralized virions that are not yet cleared by the immune system, we further analyzed subgenomic RNA (sgRNA) levels to assess active SARS-CoV-2 replication. Confirming the absence of infectious particles in the TCID_50_ assay, sgRNA levels dropped dramatically after prophylactic DZIF-10c treatment by more than three log scales in the i.n. and more than two log scales in the i.p. group (Fig. 4D). As expected, sgRNA was detected in high amounts in all control mice which indicates active replication that is prevented in case of prophylactic treatment with DZIF-10c. Again, we could observe a 6-fold stronger reduction of sgRNA in animals treated via the i.n. route compared to i.p. application.

**Fig. 4:**
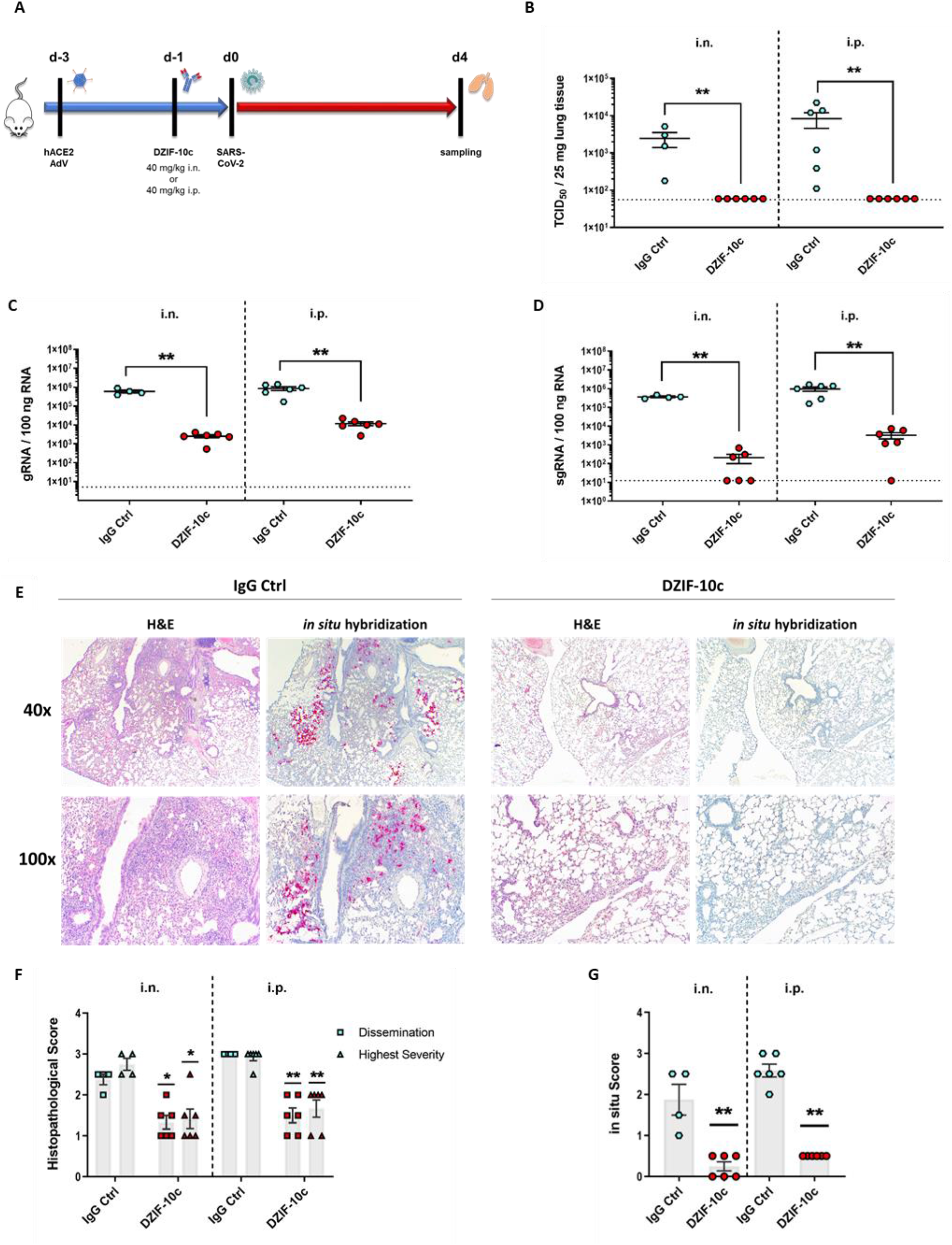
Viral load and histological analysis of hACE2-transduced BALB/c mice prophylactically treated with DZIF-10c. **(A)** Study plan of the prophylactic dose regimen. BALB/c mice were transduced with Ad_ACE2-mCherry three days before infection and treated i.n. or i.p. with 40 mg/KG body weight DZIF-10c or an IgG control antibody one day prior to challenge with SARS-CoV-2. On day four post infection, the animals were euthanized and samples were collected. Two mice from the i.n. control group had to be excluded from the analysis due to insufficient transduction efficiency. **(B)** Infectious SARS-CoV-2 titer in lung homogenates on day four post infection determined by TCID_50_ assay. **(C-D)** SARS-CoV-2 gRNA and sgRNA in lung homogenates on day four post infection determined by RT-qPCR. Error bars represent mean ± SEM. Statistical analyses were performed using Graph Pad Prism and Mann-Whitney test. *: p ≤ 0.05; **: p ≤ 0.01. Dotted lines indicate the lower limit of detection. **(E)** Histopathological analysis of the lungs by H&E staining and *in situ* hybridization of viral RNA. Images were acquired at a magnification of 40x or 100x. **(F-G)** Quantification of histopathological scores for dissemination and highest severity of inflammation as well as dissemination of viral RNA. Error bars represent mean ± SEM. Statistical analyses were performed using Graph Pad Prism and Mann-Whitney test. *: p ≤ 0.05; **: p ≤ 0.01.

Next, we evaluated the impact of DZIF-10c on lung histopathology after hematoxylin-eosin (H&E) staining. In lungs of control mice, we observed interstitial pneumonia with multiple, partially confluent foci and distinct lymphohistiocytic infiltrations in the interstitium and in the perivascular space (Fig. 4E). Importantly, prophylactic treatment with DZIF-10c markedly mitigated lung pathology with all animals showing only mild signs of inflammation. To quantify the histopathological findings, the overall dissemination of inflammation in the lung as well as the highest severity in terms of immune cell infiltration and damage to the lung architecture was assessed using a scoring system from 0 (no inflammation) to 4 (most severe) (Fig. S9). Confirming the visual observation, histopathological scores were reduced 2-fold in DZIF-10c treated mice (Fig. 4F). To compare the amounts of SARS-CoV-2 RNA, especially in inflamed areas, we further visualized viral RNA using *in situ* hybridization with SARS-CoV-2-specific probes. Consistent with our previous results, we detected high amounts of viral RNA in samples from control animals that particularly concentrated in regions with inflammation (Fig. 4E, G) whereas in DZIF-10c-treated mice, viral RNA was only detectable in single cells without spread to the surrounding tissue. This difference was more pronounced in i.n. treated mice as viral RNA was completely absent in three animals in this group. Altogether, these findings indicate that prophylactic treatment with DZIF-10c, either administered i.p. or i.n., efficiently protected hACE2-transduced mice from infection with SARS-CoV-2 as well as SARS-CoV-2-related lung pathology. Intriguingly, the i.n. administration of DZIF-10c seemed to be more effective than systemic application.

To investigate the efficiency of a therapeutic treatment with DZIF-10c, we transduced BALB/c mice intratracheally with Ad_ACE2-mCherry three days before the animals were infected intranasally with SARS-CoV-2 (Fig. 5A). In the therapeutic regimen, the mice received two doses of 40 mg/KG body weight of DZIF-10c or the isotype control antibody on days one and three after infection, either via the i.p. or the i.n. route. On day four, the animals were sacrificed and samples were taken. Clinical symptoms were absent except in one mouse of the i.n. control group, which lost 7% body weight on day four (Fig. S8). On average, 9.31×10^3^ (± 3.37 ×10^3^) TCID_50_/25 mg tissue were detected in the i.n. control group and 1.66×10^4^ (± 4.90 ×10^3^) TCID_50_/25 mg tissue in the i.p. control group (Fig. 5B). Similar to the prophylactic regimen, no infectious particles were detected in lung homogenates of DZIF-10c-treated animals indicating efficient neutralization of SARS-CoV-2 after therapeutic intervention with DZIF-10c. Interestingly, SARS-CoV-2 gRNA and sgRNA were only slightly reduced (2-3-fold) in animals receiving DZIF-10c compared to those receiving the isotype control (Fig. 5B, C). This observation was confirmed by *in situ* hybridization, which revealed a slight reduction of viral RNA in lung slices, independent on the route of antibody application (Fig. 5E, G). H&E staining revealed widespread inflammation in multiple, partially confluent foci with infiltration of lymphocytes and macrophages (Fig. 5E, F). The histopathological picture did not significantly differ between the DZIF-10c-treated and the control groups with a tendency towards a slight improvement after DZIF-10c treatment.

**Fig. 5:**
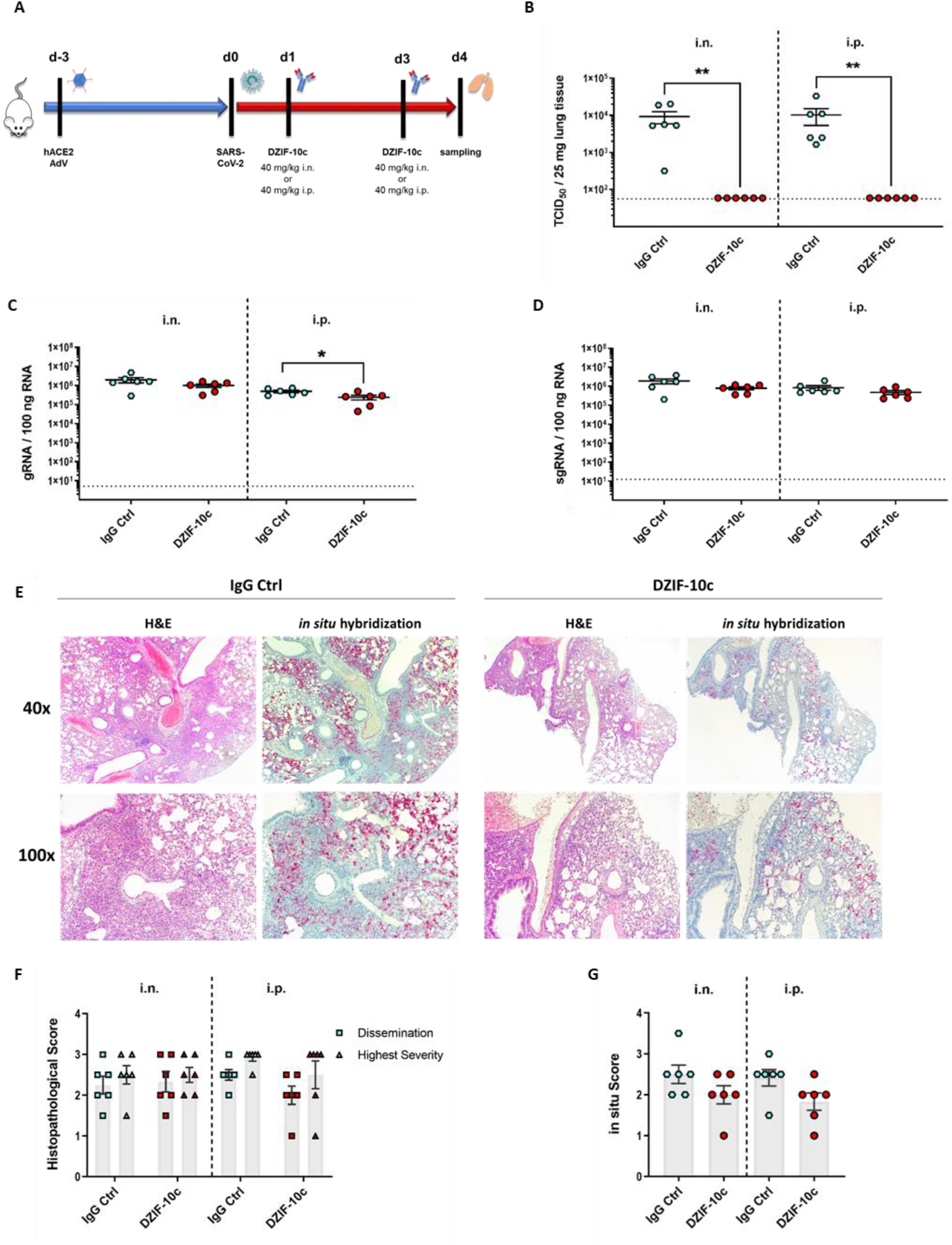
Viral load and histological analysis of hACE2-transduced BALB/c mice therapeutically treated with DZIF-10c. **(A)** Study plan of the therapeutic dose regimen. BALB/c mice were transduced with Ad_ACE2-mCherry three days before challenge with SARS-CoV-2. Mice were treated i.n. or i.p. with 40 mg/KG body weight DZIF-10c or an IgG control antibody on days one and three post infection. On day four post infection, the animals were euthanized and samples were collected. **(B)** Infectious SARS-CoV-2 titer in lung homogenates on day four post infection determined by TCID_50_ assay. **(C-D)** SARS-CoV-2 gRNA and sgRNA in lung homogenates on day four post infection determined by RT-qPCR. Error bars represent mean ± SEM. Statistical analyses were performed using Graph Pad Prism and Mann-Whitney test. *: p ≤ 0.05; **: p ≤ 0.01. Dotted lines indicate the lower limit of detection. **(E)** Histopathological analysis of the lungs by H&E staining and *in situ* hybridization of viral RNA. Images were acquired at a magnification of 40x or 100x. **(F-G)** Quantification of histopathological scores for dissemination and highest severity of inflammation as well as dissemination of viral RNA. Error bars represent mean ± SEM. Statistical analyses were performed using Graph Pad Prism and Mann-Whitney test. No statistical difference was observed in any group.

Taken together, therapeutic treatment of SARS-CoV-2 infected hACE2-transduced mice with DZIF-10c, either administered i.p. or i.n., led to a complete neutralization of infectious SARS-CoV-2 and slightly reduced amounts of viral RNA in the lungs of treated animals.

### DZIF-10c shows no evidence of enhanced infection in an *in vitro* ADE assay

Based on observations primarily made with animal models infected with SARS-CoV (29, 45), it was discussed whether ADE might play a role during SARS-CoV-2 infection. To address this question, we infected CD14^+^ human blood-derived macrophages with SARS-CoV-2 in the presence of DZIF-10c at neutralizing or non-neutralizing concentrations. To control for baseline infection and general susceptibility of our primary cell culture system, an isotype control antibody as well as infection with MERS-CoV, which is known to infect human macrophages, were included (46, 47). In addition, VeroE6 cells were included as a permissive cell line to confirm that the selected antibody concentrations were suitable to either neutralize or not neutralize the input virus. The detection of infectious MERS-CoV as well as high levels of MERS-CoV genome copies in the supernatants of both, VeroE6 cells and CD14^+^ human macrophages, confirmed that both cell types were susceptible to emerging coronaviruses (Fig. S10). As expected, neutralizing concentrations of DZIF-10c efficiently impaired infection of VeroE6 cells with SARS-CoV-2. After infection with SARS-CoV-2, significantly less viral genome copies were detected in CD14^+^ human macrophages compared to VeroE6 cells (Fig. S10A) and no infectious virus could be isolated from the supernatants at either of the tested conditions (Fig. S10B) indicating an abortive infection of the macrophages as it has been reported by others (48–50). Overall, these observations do not indicate relevant FcR-mediated enhancement of SARS-CoV-2 infection caused by DZIF-10c *in vitro*.

## Discussion

Neutralizing antibodies can be effective tools for prevention and treatment of COVID-19 (20). Our previous work identified a large panel of human monoclonal S-binding antibodies that were isolated from memory B cells from twelve convalescent COVID-19 patients (21). In the present study, we characterized one antibody of this panel (DZIF-10c) with respect to its S-binding and SARS-CoV-2-neutralizing capacity and assessed the prophylactic and therapeutic efficacy in hACE2-transduced mice. In addition, we compared systemic and topical application of DZIF-10c to investigate alternative routes of administration.

Generally, treatment efficacy of nAbs is dependent on several factors such as neutralizing titer, antibody half-life and the route of delivery (20). Our results show that DZIF-10c exerts binding and neutralization potencies comparable to antibodies that showed efficacy in clinical trials (14–16). Moreover, we analyzed the pharmacokinetics of DZIF-10c in NRG and huFcRn mice, two transgenic animal models that previously allowed reliable prediction of pharmacokinetic profiles of antiviral mAbs in humans (6, 7, 32, 34, 35). When compared to two anti-HIV-1 antibodies (3BNC117 and 10-1074), which displayed half-lives of 2-3 weeks in humans(7, 34), DZIF-10c showed similar or even prolonged stability. Exceptionally high neutralizing titers combined with a favorable pharmacokinetic profile therefore suggest DZIF-10c is suitable for clinical use.

Mechanistically, structural analysis of the DZIF-10c Fab/RBD complex by cryo-EM revealed that DZIF-10c does not directly target the ACE2 binding motif but a closely adjacent region in the prefusion conformation of SARS-CoV-2 S. In a recent publication, Barnes *et al*. proposed a classification scheme for SARS-CoV-2-RBD-specific antibodies based on binding positions and structural properties (11). Regarding its binding position, DZIF-10c belongs to class 3 binders, which are characterized by targeting a region outside of the ACE2 binding motif (Fig. S5). In contrast to the other binders in class 3, DZIF-10c is attached to the RBD only in the “up”-conformation. Due to the broad variance of resolution as evident from the local resolution map (Fig. S4C), the reconstruction in the binding region (6-10 Å) did not allow for atomic or per amino acid residue refinement of the coordinates. Therefore, the presented model represents only an approximated binding position and angle of the Fab fragment on the RBD. Further studies are necessary to define the exact epitope with atomic details.

In order to show *in vivo* efficacy of DZIF-10c, we demonstrated that both, therapeutic and prophylactic application in SARS-CoV-2-infected hACE2-transduced BALB/c mice fully neutralized infectious virus in the mouse lungs. In the prophylactic setting, the magnitude of viral load reduction as well as the reduced histopathology are in line with previous reports of other mouse transduction models (18, 42). Consistent with earlier reports from nAb-treated or vaccinated animals (13, 51), we could further observe a dramatic decrease of sgRNA compared to control mice. Since remaining gRNA levels could potentially originate from input virus or neutralized particles, sgRNA only appears during active replication (52, 53) indicating that DZIF-10c protected these animals from SARS-CoV-2 infection. After therapeutic treatment with DZIF-10c, we observed less differences in viral genome copies even though infectious virus was completely neutralized. This may be explained by the transduction mouse model that only partially reflects human SARS-CoV-2 infection (18, 54, 55). Because only successfully transduced cells are susceptible, virus spread is abolished as soon as all transduced cells have been infected. This may also be the reason why even untreated control mice challenged with SARS-CoV-2 did not develop overt clinical symptoms. Nevertheless, hACE2-transduced mice consistently display productive infection with SARS-CoV-2 indicated by high levels of viral gRNA and sgRNA, infectious progeny virus as well as interstitial pneumonia and thus represent an important and useful animal model for SARS-CoV-2.

One significant advantage of nAb immunotherapies over vaccination is their ability to provide immediate protection. However, the route of delivery can have substantial impact on the bioavailability and the clinical efficacy of a nAb. This is especially true for respiratory viruses that are predominantly present in the lung lumen. While systemic administration of nAbs leads to high antibody concentrations in serum, the availability in the lung lumen is much less pronounced, thus entailing the risk that the antibody levels at the site of action are below the effective dose (20). Several studies reported superiority of intranasal nAb delivery over systemic application in RSV or influenza A virus animal models (56–58), which is in line with a very recent study in hamsters showing efficacy of an inhaled neutralizing antibody against SARS-CoV-2 (59). Our results indicate that prophylactic treatment with DZIF-10c conveys protection against infection with SARS-CoV-2 after systemic as well as topical application. While no infectious particles could be detected in either setting, the magnitude of decrease of viral gRNA and sgRNA as well as the improvement in lung pathology were more pronounced in animals after i.n. treatment. This result indicates that DZIF-10c shows solid antiviral efficacy against SARS-CoV-2 when administered topically.

The recent emergence of SARS-CoV-2 variants with enhanced transmissibility and mutations in the RBD that potentially lead to immune escape raises concerns regarding the effectiveness of vaccines and nAbs. In particular, the single point mutations K417N, E484K and N501Y have been shown to completely abolish the neutralizing activity of several monoclonal antibodies(25–27, 60). Furthermore, a recent study demonstrated that two nAbs already in clinical use completely lost their activity against VOC B.1.351 that harbors all three of the above-mentioned substitutions (27, 28). Due to the pivotal importance of viral escape, we assessed the impact of these mutations on the neutralizing capacity of DZIF-10c. Using a pseudovirus assay, we showed that the neutralization potency of DZIF-10c was not affected by 16 out of 19 tested point mutations in the RBD, including the K417E and N501Y mutation, and retained reduced activity against pseudoviruses harboring the E484K mutation. Importantly, experiments with authentic viruses further demonstrated that DZIF-10c remained fully active against VOC B.1.1.7 and retained activity against VOC B.1.351, although with reduced potency.

When it comes to immunotherapy against viral pathogens, another area of concern is antibody-dependent enhancement of infection and disease which is based on the uptake of antibody-bound viral particles into FcR-expressing cells like macrophages or dendritic cells(45, 61). To date, there is no clear evidence that ADE plays a significant role in COVID-19 disease progression. Consistent with other studies (17, 62), we did not observe any signs of enhanced disease after DZIF-10c treatment in our mouse model (Fig. S7-S8). Furthermore, an *in vitro* ADE assay did not indicate that DZIF-10c leads to enhanced infection in FcR-bearing human macrophages. Although susceptibility of CD14^+^ human macrophages to SARS-CoV-2 may be limited, this suggests that ADE-related adverse effects of DZIF-10c are not very likely to appear in clinical trials in humans.

In summary, we characterized a novel fully human SARS-CoV-2 neutralizing antibody, DZIF-10c, which exhibited an extraordinary neutralizing potency comparable with some of the most potent anti-SARS-CoV-2 nAbs available to date. We further demonstrated that prophylactic treatment with DZIF-10c protects hACE2-transduced mice from infection with SARS-CoV-2. Moreover, our data indicate that topical administration of DZIF-10c may be a suitable delivery route that could be advantageous in clinical use. The results presented in this study made a decisive contribution to DZIF-10c having entered a phase I/II clinical trial that, for the first time, investigates an inhaled administration of a nAb targeting SARS-CoV-2 (NCT04631705).

## Materials and Methods

### ELISA analysis

ELISA plates were coated with 2 mg/ml of SARS-CoV-2 spike ectodomain, RBD, N-terminal truncated, or EBOV Makona glycoprotein (GP) ectodomain in PBS or monomeric SARS-CoV-2 spike ectodomain in 2 M Urea at 4°C overnight. Proteins were produced as previously described (21). Next day, plates were blocked with 5% BSA in PBS for 60 min at RT, incubated with primary antibody (starting concentration 10 µg/ml) in PBS for 120 min and secondary antibody (anti-human IgG-HRP; Southern Biotech 2040-05) diluted 1:2500 in 1% BSA in PBS for 60 min at RT. ELISAs were developed with ABTS solution (Thermo Fisher 002024) and absorbance was measured at 415 nm and 695 nm.

### Pseudovirus neutralization assay

SARS-CoV-2 pseudovirus particles were generated by co-transfection of individual plasmids encoding HIV Tat, HIV Gag/Pol, HIV Rev, luciferase followed by an IRES and ZsGreen, and the SARS-CoV-2 spike protein (23) into HEK 293T cells using the FuGENE 6 Transfection Reagent (Promega). Spike sequences from the following global strains REF were used: Wu01 spike (EPI_ISL_40671); BavP1 spike (EPI_ISL_406862); ARA36 spike (EPI_ISL_418432); DRC94 spike (EPI_ISL_417947); CA5 spike (EPI_ISL_408010) and NRW8 spike (EPI_ISL_414508), B.1.1.7 variant (63) and B.1.351 variant (64). Mutations were introduced by PCR into the Wu01 spike as backbone. Virus culture supernatant was harvested at 48 h and 72 h post transfection and stored at -80°C till use. The harvested virus was titrated by infecting 293T expressing ACE2 and after a 48 h incubation at 37°C and 5% CO_2_, luciferase activity was determined after addition of luciferin/lysis buffer (10 mM MgCl_2_, 0.3 mM ATP, 0.5 mM Coenzyme A, 17 mM IGEPAL (all Sigma-Aldrich), and 1 mM D-Luciferin (GoldBio) in Tris-HCL) using a microplate reader (Berthold). For neutralization assays, a virus dilution with a relative luminescence units (RLU) of approximately 1000-fold in infected cells versus non-infected cells was selected. For testing neutralization potency of DZIF-10c, serial dilutions of DZIF-10c were co-incubated with pseudovirus supernatants for 1 h at 37°C, following which 293T-ACE-2 cells were added. After a 48-hour incubation at 37°C and 5% CO_2_, luciferase activity was determined using the luciferin/lysis buffer. After subtracting background RLUs of non-infected cells, 50% inhibitory concentration (IC_50_) were calculated as the DZIF-10c concentration resulting in a 50% reduction in RLU compared to the untreated virus control wells. Each antibody dilution was tested in duplicates. IC_50_ values were calculated by plotting a dose response curve in GraphPad Prism 7.0.

### SARS-CoV-2 Virus neutralization test (VNT100)

SARS-CoV-2 neutralizing activity of human monoclonal antibodies was investigated based on a previously published protocol with slight modifications (21, 65, 66). Briefly, monoclonal antibodies were serially diluted in DMEM supplemented with 2% FBS, 1% glutamine and 1% penicillin/streptomycin (Gibco) in 96-well plates before 100 PFU SARS-CoV-2 were added to each sample. Subsequently, Vero E6 cells (Vero C1008, ATCC, Cat#CRL-1586, RRID: CVCL_0574) were washed with PBSdef, trypsinized and diluted in DMEM with 10% FBS, 1% glutamine and 1% penicillin/streptomycin. Cells were diluted in DMEM to a final concentration of 2% FBS, 1% glutamine and 1% pen/strep and 100 μl of the cell solutions was added to virus/antibody samples, corresponding to approximately 20.000 cells/well. Neutralization was defined as absence of cytopathic effect compared to virus controls (IC_100_). The following controls were included: back titration of virus dilution, positive control as an inter-assay neutralization standard (cells infected with SARS-CoV-2 and treated with an antibody with a known neutralizing titer), negative control (cells without infection and antibodies), cytotoxicity control (cells without infection, only treated with antibodies).

### Cryo-electron microscopy

Fab fragments were generated and purified from full length IgG using a Pierce™ Fab Preparation Kit (Thermo Fisher). Purified Fabs were mixed with SARS-CoV-2 S protein, Super stable trimer (AcroBiosystems) (1.1:1 molar ratio Fab per protomer) to a final protein concentration of 0.2 mg/ml and incubated on ice for 30 min. 3.5 µl of the complex solution were deposited onto a C-flat 1.2/1.3-3C holey carbon copper grid (Electron Microscopy Sciences) that had been freshly glow-discharged for 1 min at 20 mA using a PELCO easiGLOW (Ted Pella). Samples were vitrified in 100% liquid ethane using a Leica EM GP2 automatic plunge freezer (Leica Microsystems) after blotting at 10 °C and 85% humidity for 4 s. cryo-EM images were collected on a Titan Krios transmission electron microscope (Thermo Fisher) at 300 kV using a K3 detector (Gatan) in super-resolution counting mode. Images were energy filtered (20 eV slit) and collected automatically using EPU v. 1.2 software (Thermo Fisher). Each image was composed of 50 individual frames with a total exposure dose of 50 e^-^/Å^2^ and a pixel size of 0.415 Å. Single particle data processing was performed in cryoSPARC v2.15 (Structura Biotechnology Inc.) as described below. Super resolution movies were patch motion corrected, Fourier-cropped (factor 1/2) and dose weighted before estimating CTF parameters using the Patch CTF job type. Particles were picked using the reference-free Blob picker, extracted with a box size of 360 pixels (0.83 Å/pixel) and subjected to 2D classification. The best class averages were selected manually (226k particles) to create an ab initio 3D model followed by a homogeneous refinement. Particles were then 3D-classified into four classes and the best resolved classes were selected for a final round of homogenous and subsequent non-uniform refinement (142k particles). For interpretation of the reconstructed 3D cryo-EM map of the Fab-S protein complex PDB 6VSB (S protein) and PDB 7C01 were used (Fab fragment). The Fab fragment coordinates of PDB 7C01 were edited with Sculptor in Phenix (Lawrence Berkeley National Laboratory) to match the sequence of DZIF-10c Fab. Atomic coordinates were rigid body fitted into the 3D cryo-EM map using UCSF Chimera v1.13.1. The resolution of the map did not allow for atomic or per amino acid residue refinement of the coordinates. All Figures were created in UCSF Chimera v1.13.1.

### Tissue culture infectious dose 50 (TCID_50_) Assay

The amount of infectious SARS-CoV-2 particles from cell culture supernatants or lung homogenates was determined by TCID_50_ assay. Vero E6 cells were cultured in DMEM with 2% FBS, 1% glutamine, 1% penicillin/streptomycin in 96-well plates and were inoculated with 10-fold serial dilutions of supernatant samples or 5-fold serial dilutions of lung homogenates. At four days post infection, SARS-CoV-2 CPE was evaluated, and titers per ml or 25 mg lung tissue were calculated using the Spearman-Kaerber method (67).

### Detection of genomic RNA (gRNA) by quantitative real-time reverse transcription PCR (RT-qPCR)

In order to quantify viral gRNA, nucleic acids were isolated from cell lysates or lung homogenates using the RNeasy Mini Kit (Qiagen) according to manufacturer’s instructions. Total RNA amount was measured using a NanoDrop ND-100 spectrophotometer.

For analysis of SARS-CoV-2 genome copies, RNA was reverse transcribed and viral genome copies quantified by real-time PCR using the OneStep RT-PCR Kit (Qiagen) and the StepOne Real-Time PCR System (Applied Biosystems). Primers and probes targeting the E gene of SARS-CoV-2 (E Assay_First Line Screening) as well as a positive control plasmid were purchased from idtdna (Berlin, Germany) (68). Reverse transcription and amplification were performed using the following protocol: 55°C for 30 min, 95°C for 15 min followed by 45 cycles of 95°C for 5 s, 60°C for 15 s and 72°C for 15 s. Quantification was carried out using a standard curve based on 10-fold serial dilutions of a plasmid DNA comprising the target region ranging from 10^3^ to 10^6^ copies.

E_Sarbeco_F1: ACAGGTACGTTAATAGTTAATAGCGT

E_Sarbeco_R2: ATATTGCAGCAGTACGCACACA

E_Sarbeco_P1: FAM-ACACTAGCCATCCTTACTGCGCTTCG-BBQ

MERS-CoV genome copies were determined using a previously published protocol (40, 69). Briefly, RNA was reverse transcribed and viral genome copies quantified by real-time PCR using the SuperScript III OneStep RT-PCR Kit (Invitrogen Life Technologies) and the StepOne Real-Time PCR System (Applied Biosystems). Primers and probes targeting the E gene of MERS-CoV were purchased from Tib-Molbiol (Germany). Reverse transcription and amplification were performed using the following protocol: 55°C for 20 min, 95°C for 3 min followed by 45 cycles of 94°C for 15 s, 58°C for 30 s. Quantification was carried out using a standard curve based on 10-fold serial dilutions of appropriate cloned RNA ranging from 10^1^ to 10^6^ copies.

upE_Fwd: GCAACGCGCGATTCAGTT

upE_Rev: GCCTCTACACGGGACCCATA

upE_Prb: FAM-CTCTTCACATAATCGCCCCGAGCTCG-TAMRA

### Detection of subgenomic RNA (sgRNA) by quantitative real-time reverse transcription PCR (RT-qPCR)

Subgenomic RNA was determined according to a previously published protocol (13, 52). Nucleic acids were isolated as described before. SARS-CoV-2 subgenomic RNA of the E gene was reverse transcribed and copy numbers quantified by real-time PCR using the SuperScript III OneStep RT-PCR Kit (Invitrogen Life Technologies) and the StepOne Real-Time PCR System (Applied Biosystems). Primers and probes targeting the leader and the E gene of SARS-CoV-2, respectively, were purchased from Tib-Molbiol (Germany). Reverse transcription and amplification were performed using the following protocol: 55°C for 20 min, 95°C for 3 min followed by 45 cycles of 95°C for 10 s, 56°C for 15 s and 72°C for 15 s. Quantification was carried out using a standard curve based on 10-fold serial dilutions of a plasmid DNA comprising the target region ranging from 10^1^ to 10^7^ copies.

sgLead_SARS2_F: CGATCTCTTGTAGATCTGTTCTC

sgE_SARS2_R: ATATTGCAGCAGTACGCACACA

sgE_SARS2_P: FAM-ACACTAGCCATCCTTACTGCGCTTCG-BBQ

### Pharmacokinetic Profile of DZIF-10c in NRG and FcRn Mice

NOD.Cg-*Rag1*^*tm1Mom*^ *Il2rg*^*tm1Wjl*^/SzJ (NRG) and B6.Cg-*Fcgrt*^*tm1Dcr*^ *Prkdc*^*scid*^ Tg(FCGRT)32Dcr/DcrJ (FcRn) mice (The Jackson Laboratory) were bred and maintained at the Decentralized Animal Husbandry Network of the University of Cologne, and the experiments were authorized by the State Agency for Nature, Environmental Protection, and Consumer Protection North Rhine-Westphalia (84-02.04.2015.A353). To determine *in vivo* pharmacokinetic profiles of individual antibodies in NRG and FcRn mice, longitudinal serum samples were collected after a single intravenous injection of 0.5 mg of monoclonal antibody in PBS. Serum samples were stored at -20°C until analysis and a sample obtained from each mouse before the start of the experiment was used to confirm the baseline absence of human IgG. Human IgG serum concentrations were determined as described previously with minor modifications (33). High-binding ELISA plates (Corning) were coated with goat anti-human IgG (Jackson ImmunoResearch) at a concentration of 2.5 µg/ml for 10 h at room temperature (RT, NRG mice) or 2 h at 37°C (FcRn mice), followed by blocking with blocking buffer (2% BSA (Carl Roth), 1 µM EDTA (Thermo Fisher), and 0.1% Tween-20 (Carl Roth) in PBS) for 80 min at 37°C (NRG mice) or 120 min at RT (FcRn mice). Subsequently, a human IgG1 kappa standard purified from myeloma plasma (in duplicates per plate, Sigma-Aldrich) and serum samples (starting at a 1:20 dilution) were incubated in serial dilutions in PBS for 75-90 min at room temperature (RT). For detection, HRP-conjugated goat anti-human IgG (Jackson ImmunoResearch) diluted 1:1,000 in blocking buffer was applied for 75-120 min at RT. Finally, optical density at 415 nm was determined using a microplate reader (Tecan) after addition of ABTS (Thermo Fisher). Between each step, plates were washed with 0.05% Tween-20 in PBS. Human serum IgG concentrations were calculated using the plate-specific IgG1 standard curve.

### Analysis of DZIF-10c *in vivo* efficacy after SARS-CoV-2 challenge

All challenge animal experiments were performed in accordance with the German animal protection laws and were authorized by the regional authorities (RP Gießen, G44/2020). The hACE2 transduction mouse model for SARS-CoV-2 and associated procedures were designed based on a previously established model for MERS-CoV (39, 40, 70) and were described recently (41). Briefly, 6-8 week old BALB/c mice were purchased from Charles River Laboratories and housed under specific pathogen-free conditions in isocages in the animal facility at the Institute of Virology Marburg. Prior to viral challenge, all mice were inoculated intratracheally with 5×10^8^ PFU Ad_ACE2-mCherry-mCherry (cloned at ViraQuest Inc., North Liberty, IA, USA) under short ketamine/xylazine anesthesia in order to induce pulmonary expression of hACE2. Three days post transduction, mice were inoculated via the intranasal route with 1.5×10^4^ TCID_50_ SARS-CoV-2 (BavPat1/2020 isolate, European Virus Archive Global # 026V-03883) in the BSL-4 facility (Institute of Virology, Philipps University Marburg, Germany). Mice were monitored daily and clinical scores including body weight changes were documented. On day four post infection mice were sacrificed and lung samples were collected. Lung samples were taken from the upper left lung lobe and were homogenized in 1 ml DMEM with ceramic and glass beads (Lysing Matrix H 500, 2 ml tube, MP Biomedicals) in a mixer mill (Retsch Schwingmühle MM 400) for 5 min at 30 Hz. To remove tissue debris, homogenates were centrifuged for 5 min at 2,400 rpm. 40 µl of fresh homogenates were used for a TCID_50_ assay. Further 100 µl of the homogenate were used for RNA isolation.

Application of monoclonal antibodies (DZIF-10c or IgG isotype control) was performed via the intranasal or the intraperitoneal route at a dose of 40 mg/KG. Antibody treatment was conducted on day one before infection (prophylactic regimen) or twice on day one and three after infection (therapeutic regimen). In order to ensure a safe administration of antibodies, mice were anesthetized shortly with isoflurane before each treatment.

### Histopathological examination of lung tissue

Lungs were collected on day four post challenge with SARS-CoV-2 and processed for histological analysis as described before (40). Tissue was fixed in formalin and embedded in paraffin. For histopathological analysis, sections were cut with a Leica RM2255 microtome (Leica Biosystems) and stained with hematoxylin and eosin (H&E). To investigate the presence of viral RNA, lungs were mounted on slides and analyzed via *in-situ* hybridization. To this end, the RNAscope® 2.5 HD Assay – RED Kit from Bio-Techne (Cat. No. 322360) was used according to the manufacturer’s instructions. Briefly, mounted slides were baked at 60°C, deparaffinized with xylene and 100% ethanol and pretreated with RNAscope® Pretreatment Reagents (Cat. No. 322300 and 322000) to enable access to the target RNA. Subsequently, a RNA-specific probe, targeted against the S gene of the SARS-CoV-2 (Cat. No. 848561), was hybridized to the RNA. Fast Red substrate was administered to the samples, allowing signal detection. The slides were counterstained with Gill’s Hematoxylin I and 0.02 % ammonia water. A RNAscope® Negative Control Probe (Cat. No. 310043) was used in parallel to monitor background staining.

### Statistical analysis

In order to test for statistical differences in viral load analyses (RT-qPCR, TCID_50_ assays) and histopathological scores between treated and non-treated animals, a nonparametric analysis using the Mann-Whitney test was performed. Results were declared significant at p < 0.05. All statistical analyses were performed using GraphPad Prism.

## List of Supplementary Materials

### Supplementary Methods

Fig. S1: Binding of DZIF-10c to different forms of SARS-CoV-2 S

Fig. S2: Affinity analysis of DZIF-10c and SARS-CoV-2 S RBD-HIS measured by SPR.

Fig. S3: Neutralization of SARS-CoV-2 pseudoviruses bearing different mutations in the spike RBD

Fig. S4: Resolution of the cryo EM reconstruction

Fig. S5: Classification scheme for SARS CoV 2 RBD specific antibodies according to Barnes et al

Fig. S6: RT-qPCR-based analysis of mCherry levels in lung homogenates of hACE2-transduced mice

Fig. S7: Body Weight and Clinical Scores of prophylactically treated SARS-CoV-2 infected mice

Fig. S8: Body Weight and Clinical Scores of therapeutically treated SARS-CoV-2 infected mice

Fig. S9: Representative examples for histopathology scoring

Fig. S10: *In vitro* ADE Assay

## Acknowledgments

We gratefully thank Wilhelm Bertrams, Anna Lena Jung and Bernd Schmeck for kindly providing human blood-derived macrophages and Stefan Pöhlmann for providing the Vero76-TMPRSS2 cells. We further gratefully acknowledge Torsten Hain, Jan Philipp Mengel and Nadine Biedenkopf for the help with sequencing of virus isolates and Tobias Nolden for his support with the cloning of expression plasmids for the pseudovirus variants. We acknowledge Christian Keller and the European Virus Archive Global (EVAg) for providing virus isolates used in this study (details in methods section). Moreover, we thank Astrid Herwig, Susanne Berghöfer, Jana Schneider, Dirk Becker, Martina Huxol, Gotthard Ludwig, Michael Schmidt, Nikolai Prill and Julia Eppensteiner for excellent technical support.

## Data and materials availability

All relevant data are within the manuscript and its Supporting Information files. Materials/samples used in the analysis described in this manuscript may be made available to qualified, academic, noncommercial researchers through a material transfer agreement upon request.

