## Supplementary Materials for "Intranasal administration of a monoclonal neutralizing antibody protects mice against SARS-CoV-2 infection"

*Corresponding author:

**This PDF file includes:**

Supplementary text (Methods)
Figures S1 to S10
Legends for Datasets S1 to S10

### Supplementary Methods

#### Cell culture

Vero E6 cells (Vero C1008, ATCC, Cat#CRL-1586, RRID: CVCL_0574) were maintained in Dulbecco`s Modified Eagle`s Medium (DMEM) supplemented with 10% FBS, 1% Glutamine and 1% penicillin/streptomycin (Gibco). Virus infections were performed in DMEM with 2% FBS, 1% Glutamine and 1% penicillin/streptomycin.

Cultivation and differentiation of human blood-derived macrophages was performed at the Institute for Lung Research (Marburg, Germany) as described previously. Briefly, macrophages were obtained from primary monocytes isolated by MACS CD14 positive selection from healthy donor buffy coats provided by the Centre for Transfusion Medicine and Haemotherapy, University Hospital Giessen and Marburg (Germany). All blood donors gave informed written consent for use of their blood samples for scientific purposes (Ethics approval number: 161/17). Freshly isolated monocytes were seeded in ultra-low attachment plates (Corning) and were left to adhere for 2 h in supplement-free RPMI medium (Gibco). After adhesion of cells, differentiation was initiated by addition of 1% human AB serum (Sigma) and monocytes were cultivated for seven days. Maturation to blood-derived macrophages was confirmed by microscopy. Cells were detached by incubation with pre-warmed PBS for 10 min and then seeded at a density of $3x10^5$ cells/12well in 1 mL of fresh RPMI media. All cells were cultivated at 37°C and 5% $CO_2$.

#### Viruses

All work with infectious SARS-CoV-2 was performed in the BSL-4 facility at the Institute of Virology, Philipps University Marburg (Marburg, Germany). BavPat1/2020 isolate (#026V-03883) was purchased from European Virus Archive Global (EVAg). The B.1.1.7 variant (BioProject no. PRJNA721582) was a clinical isolate and was isolated at the Institute of Virology, Philipps University Marburg (Marburg, Germany). The B.1.351 (GenBank Accession no. MW822592) and P.2 (GenBank Accession no. MW822593) variants were clinical isolates and were isolated at the Institute for Medical Virology, University Hospital Frankfurt

(Goethe University Frankfurt am Main, Germany). All viruses were propagated on Vero76-TMPRSS2 cells using DMEM supplemented with 2% FBS, 1% Glutamine and 1% penicillin/streptomycin (Gibco). On day two (BavPat1/B.1.1.7/B.1351) or three (P.2) after infection, cell culture supernatants were harvested, and titers were determined by immunoplaque and $TCID_{50}$ assay on VeroE6 cells.

**Production of DZIF-10c**

Original V-region sequences of the antibody HbnC3t1p1_F4 (22) were formatted unaltered onto human IgG1 backbone (G1m3 allotype) with the C-terminal lysine removed. Light and heavy chains were configured onto separate pTT5 (licensed from National Research Council of Canada) expression vectors. Briefly, V-regions were codon-optimized for mammalian expression, ordered as vector-overlapping dsDNA fragments from idtdna (Berlin, Germany), and cloned into light and heavy chain pTT5 vectors by Infusion reaction methodology (Clontech). Standard transformation procedure was completed utilizing Stellar cells (Clontech) and 500 ml *E. coli* cultures (LB media with 100 mg/ml carbenicillin) were grown to generate substantial amounts of plasmid DNA for Megaprep plasmid plus purification (Qiagen). Finalized plasmid DNA was sequenced externally (Genewiz) and matched against reference sequence using Lasergene Seqman software. Following plasmid preparation, all antibodies (DZIF-10c or anti-TNP IgG Isotype control) were expressed in CHO-3E7 (CHO-E) cells using previously described protocols. Briefly, cells are maintained in an actively dividing state in FreeStyle CHO (FS-CHO) medium before transfection with TransIT Pro (Mirus Bio) following manufacturer's recommendations. The transfected culture was maintained for ten days, and harvest was done by centrifuging and sterile filtration. Then, cell culture supernatants were loaded onto MabSelect SuRe column (Cytiva, product number 11003494) pre-equilibrated with Dulbecco's phosphate buffered saline (DPBS). The columns were then washed with DPBS, DPBS plus 1.0 M NaCl and then DPBS. Then the bound proteins were eluted from the columns with 30 mM sodium acetate (pH 3.5) and the pools was neutralized with 1% volume to volume of 3 M sodium acetate (pH ≈ 9). The neutralized samples were then sterilely filtered with filtration units, followed by measurements of protein concentration, endotoxin level and purify check by SDS-PAGE as well as aSEC. The impurities (*i.e.,* aggregates) were then further removed with CEX chromatography by loading the sample onto prepacked POROS HS50 column (Thermo Scientific, A36637), washed and then eluted with a salt gradient. The fractions of the eluate were analyzed by aSEC. The high percent monomer fractions were pooled together, and salt concentration was adjusted to 100 mM NaCl. The proteins were then sterilely filtered and final quality and quantity were assessed (i.e., protein concentration, endotoxin level, percent monomer by aSEC).

**Surface Plasmon Resonance (SPR)**

The SPR assays were performed using a Biacore 8K system and a CM5 sensor chip (GE Healthcare). The running buffer for this experiment and all dilutions were done in 1 X HBS-EP+ (GE Healthcare Life Sciences BR100669). The CM5 sensor chip was activated with equal mixture of EDC/NHS for 420 sec at a flow rate of 10 ml/min and immobilized with Protein A/G (50 mg/ml in 10 mM acetate pH 4.5) for 420 sec at a flowrate of 10 ml/min resulting in ≈ 2400-2800 RU of Protein A/G on the surface. Subsequently, the sensor chip was deactivated with 1M ethanolamine HCl for 420 sec at a flowrate of 10 ml/min. DZIF-10c (1 mg/ml) was captured on the Protein A/G surface for 60 sec at a flowrate of 10 ml/min resulting in capture levels of ≈ 150 RU. The analyte (SARS-COV2 RBD-His) was injected over the captured ligand for 120 sec at a flowrate of 30 ml/min. The dissociation was done for 600 sec. The concentrations of the analyte were as follows: 0 nM, 1.56 nM, 3.13 nM, 6.25 nM, 12.5 nM, and 25 nM. After each analyte injection was complete, the surface was regenerated by injecting 0.85% phosphoric acid for 30 sec at a flowrate of 30 ml/min. The analyte interaction with sensor surface (flow cell 1) and blank (HBS-EP+ or 0 nM analyte) were subtracted from the raw data. Sensorgrams were then fit globally to 1:1 Langmuir binding to provide on-rate ($k_a$), off-rate ($k_d$), and affinity ($K_D$) values. The binding experiments for SARS-COV-2 RBD-His were performed three separate times using fresh dilutions.

***In vitro* ADE Assay**

For assessment of possible DZIF-10c-related enhancement of SARS-CoV-2 infection (ADE), human blood-derived macrophages were infected with SARS-CoV-2 in presence of different concentrations of DZIF-10c or IgG isotype control antibodies. $2.5x10^5$ macrophages were seeded in 12 well plates in 1 ml RPMI medium supplemented with 1% Glutamine, penicillin/streptomycin and non-essential amino acids. After adhesion, 10% FBS was added and the cells were incubated for three to four days. Prior to infection with SARS-CoV-2 non-neutralizing IgG control antibodies or DZIF-10c in neutralizing and sub-neutralizing concentrations, respectively, were incubated for 1 h at 37°C together with 800 50% tissue culture infective doses ($TCID_{50}$) SARS-CoV-2 (BavPat1/2020 isolate, European Virus Archive Global # 026V-03883). Directly before inoculation of macrophages with the antibody/virus mixture, 500 µl of the macrophage cell culture medium was transferred to a new 12 well plate and mixed with fresh supplemented 10% RPMI medium (storage medium). Subsequently, 800 µl of the antibody/virus mixture was added to the macrophages and cells were incubated for 4 h at 37°C. Afterwards, antibody/virus-containing medium was discarded and 1 ml storage medium was added to the cells. At four days post infection, CPE was evaluated and supernatants were collected and stored at -80°C. RNA isolation from cell lysates was performed using the RNeasy Mini Kit (Qiagen) according to manufacturer's instructions. In control settings, the experimental protocol was modified regarding virus and cells. MERS-CoV (EMC/2012) was used to show general susceptibility of macrophages to infection with coronaviruses. Vero E6 cells were used to demonstrate infectivity of the SARS-CoV-2 isolate as well as the neutralizing activity of DZIF-10c in this experimental setting.

**Detection of mCherry mRNA by quantitative real-time reverse transcription PCR (RT-qPCR)**

To confirm transduction with AdV-hACE2, mRNA levels of the reporter mCherry was determined according to a previously published protocol (40). Nucleic acids were isolated from lung homogenates as described before. mCherry mRNA was reverse transcribed and copy numbers quantified by real-time PCR using the SuperScript III OneStep RT-PCR Kit (Invitrogen Life Technologies) and the StepOne Real-Time PCR System

(Applied Biosystems). Primers and probes targeting the mCherry gene were purchased from Tib-Molbiol (Germany). Reverse transcription and amplification were performed using the following protocol: 50°C for 30 min, 95°C for 15 min followed by 40 cycles of 95°C for 15 s, 48°C for 30 s and 72°C for 20 s. Quantification was carried out using a standard curve based on 10-fold serial dilutions of appropriate cloned RNA ranging from $5x10^2$ to $5x10^5$ copies.

mcherry181for: CATGGTAACGATGAGTTAG

mcherry287rev: GTTGCCTTCCTAATAAGG

mcherry probe: FAM- TACCACCTTACTTCCACCAATCGG-BBQ

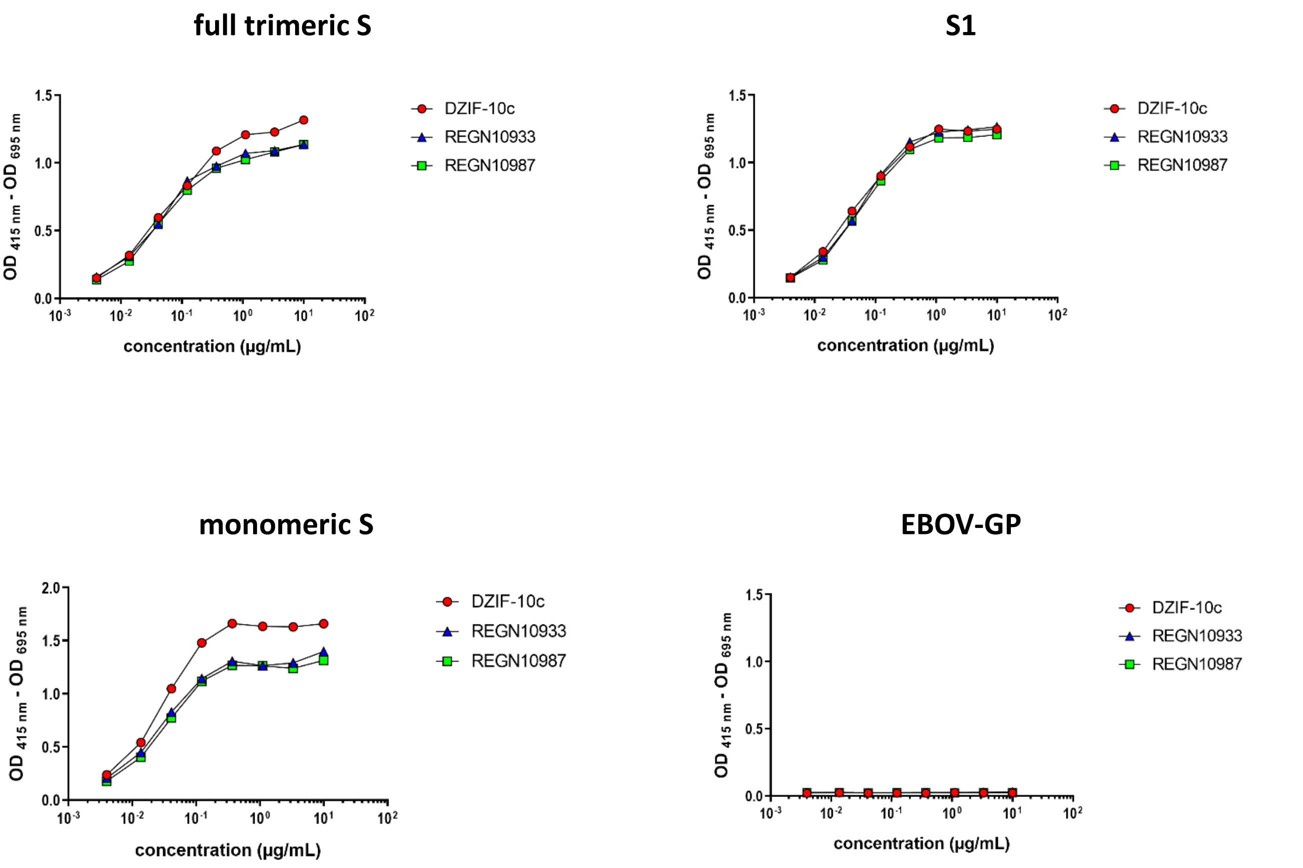


**Fig. S1: Binding of DZIF-10c to different forms of SARS-CoV-2 S**

Interaction of DZIF-10c with full trimeric S **(A)**, S1 **(B)**, monomeric S **(C)** and EBOV-GP **(D)** measured by ELISA.

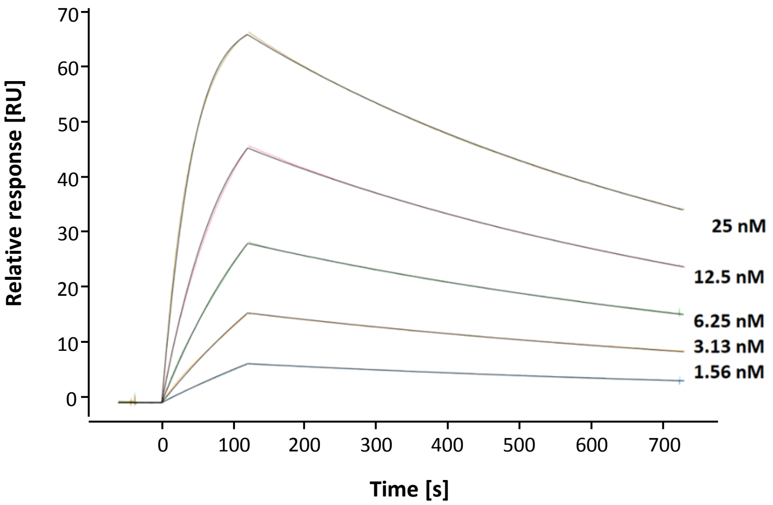


**Fig. S2: Affinity analysis of DZIF-10c and SARS-CoV-2 S RBD-HIS measured by SPR.**

DZIF-10c (1 µg/ml) was captured on the Protein A/G surface and the analyte was injected over the captured ligand. The concentrations of the analyte (SARS-COV-2 RBD-His) were as follows: 0 nM, 1.56 nM, 3.13 nM, 6.25 nM, 12.5 nM, and 25 nM. The analyte interaction with sensor surface (flow cell 1) and blank (HBS-EP+ or 0 nM analyte) were subtracted from the raw data. Sensorgrams were then fit globally to 1:1 Langmuir binding to provide on-rate (ka), off-rate (kd), and affinity (KD) values. The diagram shows a representative SPR sensorgram from three experiments.

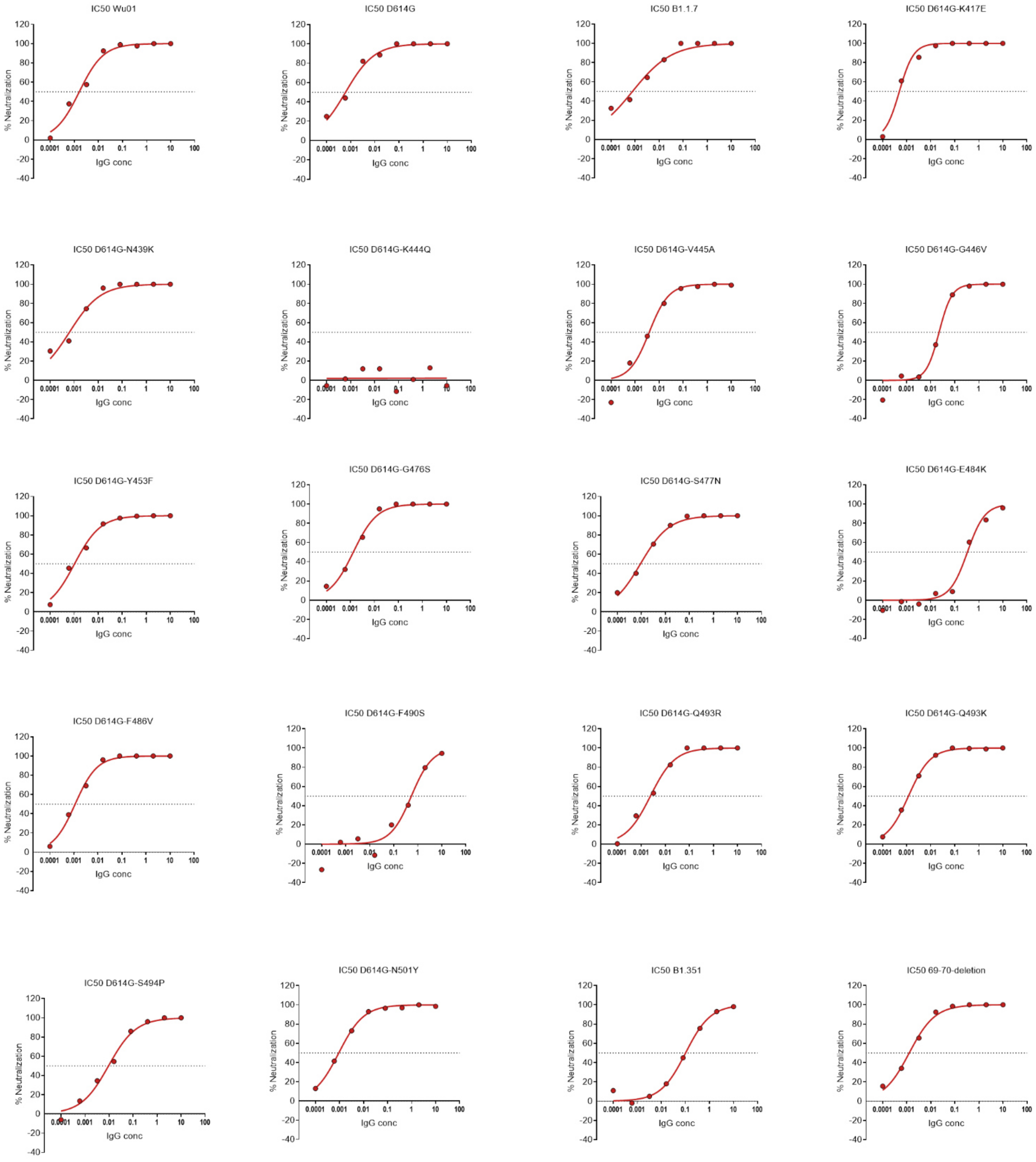


**Fig. S3: Neutralization of SARS-CoV-2 pseudoviruses bearing different mutations in the spike RBD**

Neutralizing activity of DZIF-10c against SARS-CoV-2 pseudoviruses bearing S proteins with point mutations found in circulating variants of concern. The dotted line indicates 50% neutralization ($IC_{50}$).

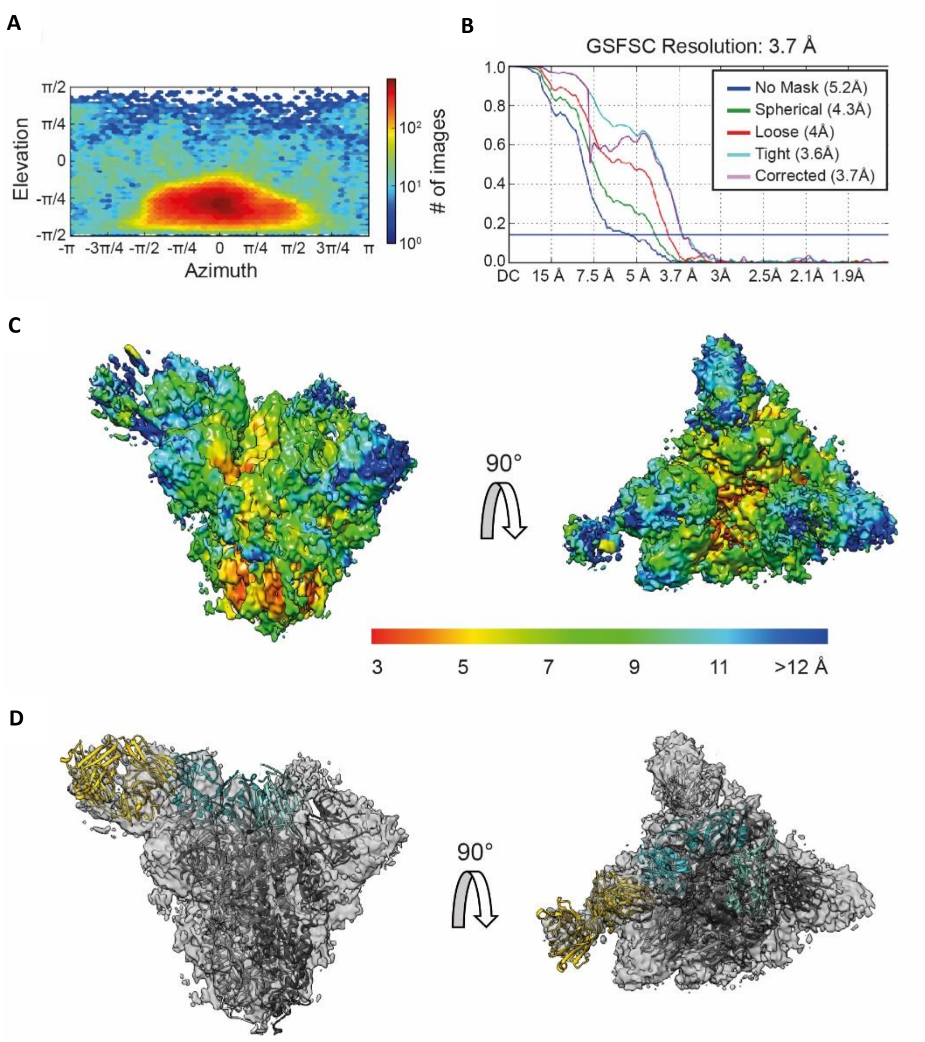


**Figure S4: Resolution of the Cryo-EM reconstruction.**

**(A)** Viewing Direction Distribution heat map, showing the predominance of top views. **(B)** Gold standard FSC curve. The 0.143 criterion reports a global resolution of 3.7 Å. **(C)** Local resolution map showing the variance of resolution in the reconstruction of the complex using a rainbow color code from red (high resolution) to blue (low resolution). Left: side view; Right: top view. **(D)** Rigid body fits of PDB models 6VSB (S protein protomers: grey shades; RBDs: cyan shades) and 7C01 (Fab fragment, yellow) overlaid with the reconstructed, transparent cryo-EM map (grey). Left: side view; Right: top view.

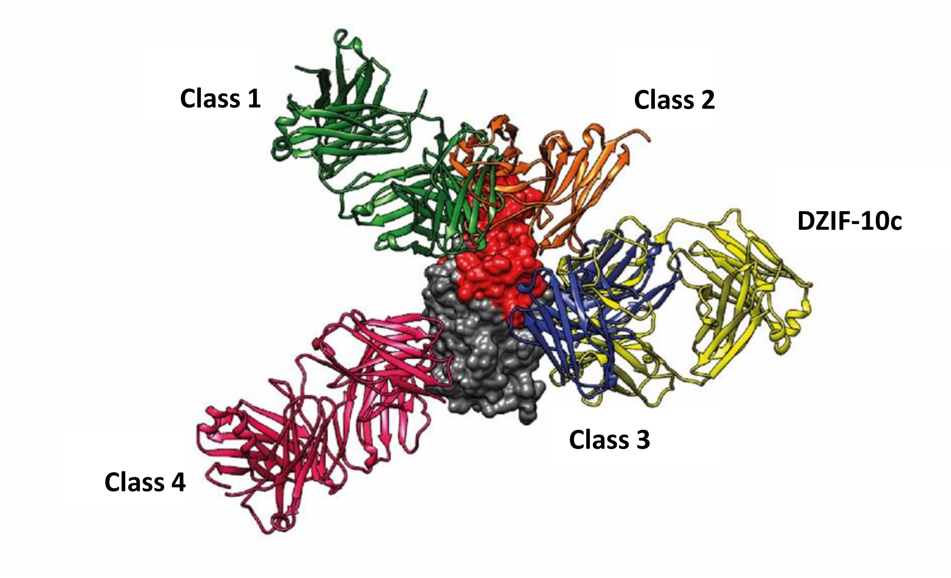


**Figure S5: Classification scheme for SARS-CoV-2-RBD-specific antibodies according to Barnes et al. *(11)*.** Based on the binding position, DZIF-10 belongs to class 3 binders. Grey: Spike RBD; Red: ACE-2 binding motif.

A

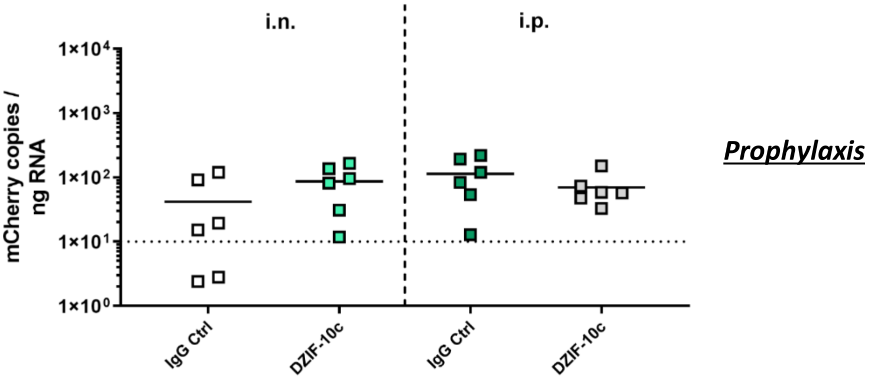


B

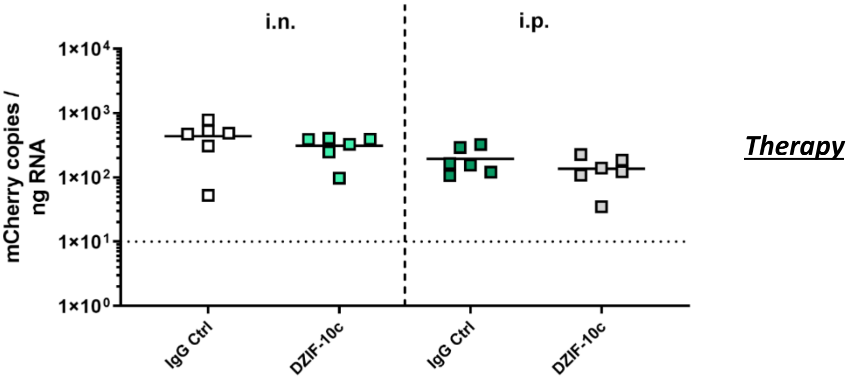


**Fig. S6: RT-qPCR-based analysis of mCherry levels in lung homogenates of hACE2-transduced mice**

BALB/c mice were transduced with AdV-hACE2 encoding for the reporter mCherry three days before infection. Mice were treated i.n. or i.p. with 40 mg/KG body weight DZIF-10c or an IgG control antibody on day one before **(A)** or on day one and three after **(B)** challenge with SARS-CoV-2. On day four post infection, the animals were euthanized and samples were collected. Squares indicate mCherry mRNA copies in lung homogenates on day four post infection determined by RT-qPCR. The dotted lines show the limit for inclusion at 10 mCherry copies / ng RNA. Two mice from the i.n. control group in the prophylactic study (A) had to be excluded from the analysis due to insufficient transduction efficiency indicated by mCherry copy numbers below 10 copies / ng RNA.

A

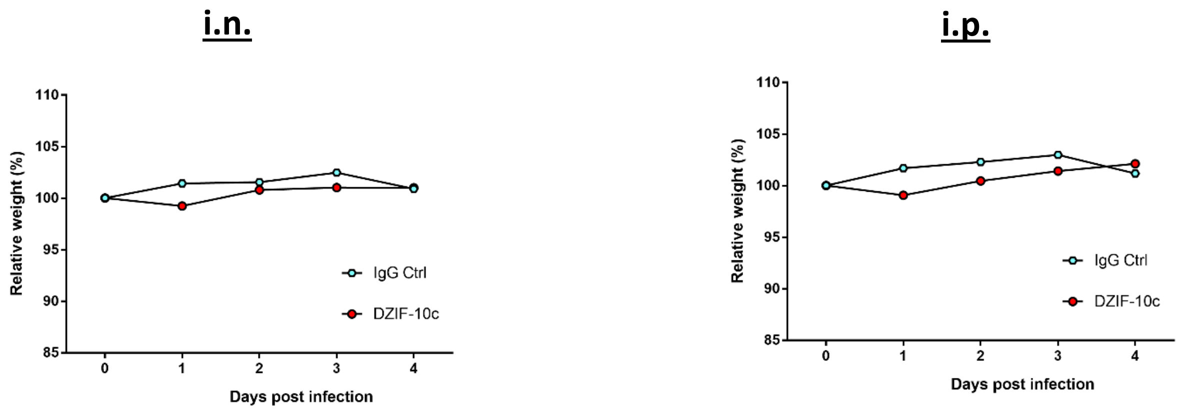


B

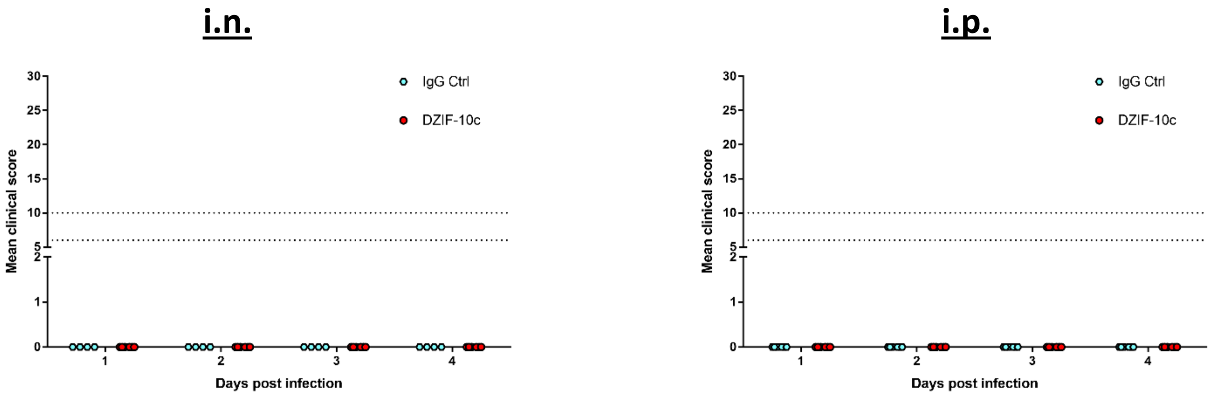


**Fig. S7: Body Weight and Clinical Scores of prophylactically treated SARS-CoV-2 infected mice**

**(A)** Changes in body weight for i.n. or i.p. treated mice after SARS-CoV-2 challenge. Dots depict group means at the respective time point. **(B)** Clinical scores for i.n. or i.p. treated mice after SARS-CoV-2 challenge. Clinical scores are composed of body weight, spontaneous behavior and general condition. Dotted lines represent clinical end points. Animals are euthanized at a score of 10 or of 6 on two consecutive days. Dots depict individual scores for each animal at the respective time point.

A

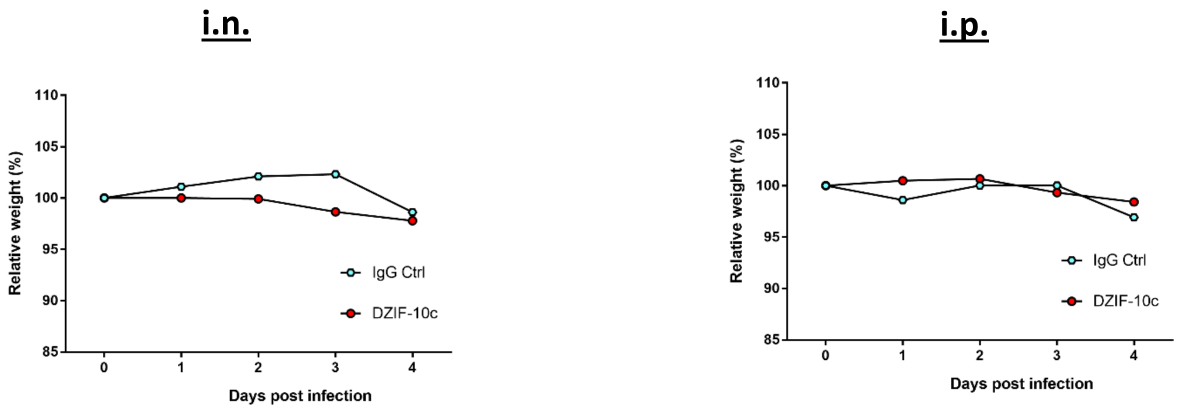


B

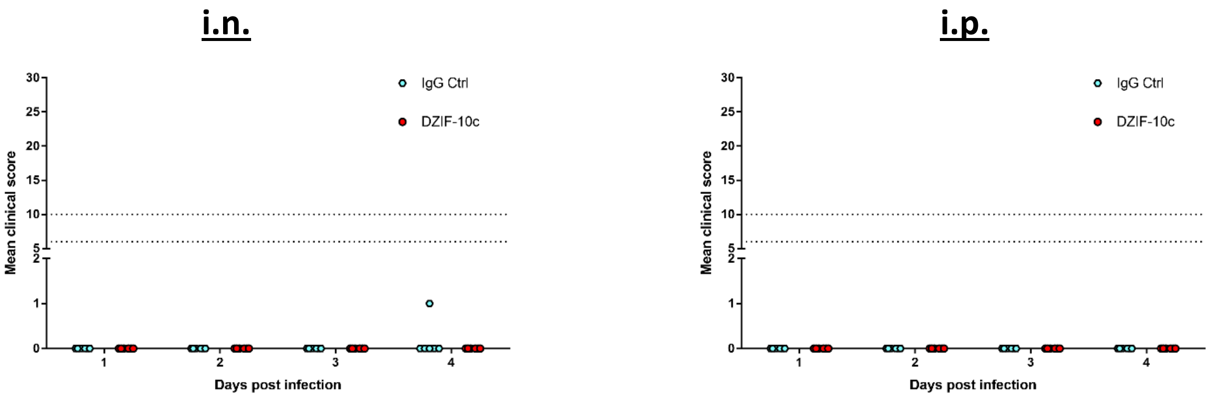


**Fig. S8: Body Weight and Clinical Scores of therapeutically treated SARS-CoV-2 infected mice**

**(A)** Changes in body weight for i.n. or i.p. treated mice after SARS-CoV-2 challenge. Dots depict group means at the respective time point. **(B)** Clinical scores for i.n. or i.p. treated mice after SARS-CoV-2 challenge. Clinical scores are composed of body weight, spontaneous behavior and general condition. Dotted lines represent clinical end points. Animals are euthanized at a score of 10 or of 6 on two consecutive days. Dots depict individual scores for each animal at the respective time point.

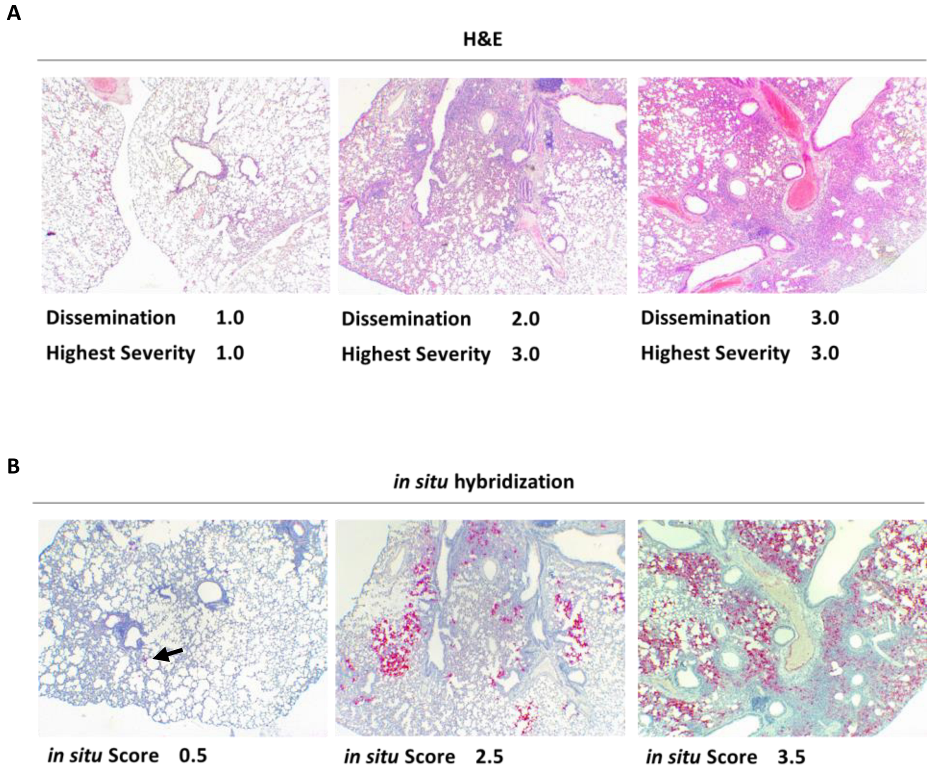


**Fig. S9: Representative examples for histopathology scoring**

Histopathological analysis of the lungs by H&E staining **(A)** and *in situ* hybridization of viral RNA **(B)**. Images were acquired at a magnification of 40x. Histopathological scores of the respective samples are specified below. The black arrow in Panel B highlights one single spot where viral RNA was detected.

A

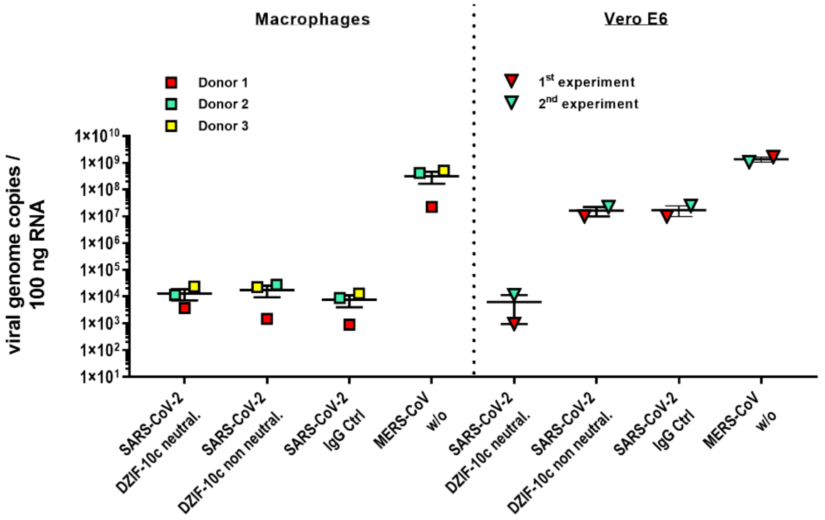


B

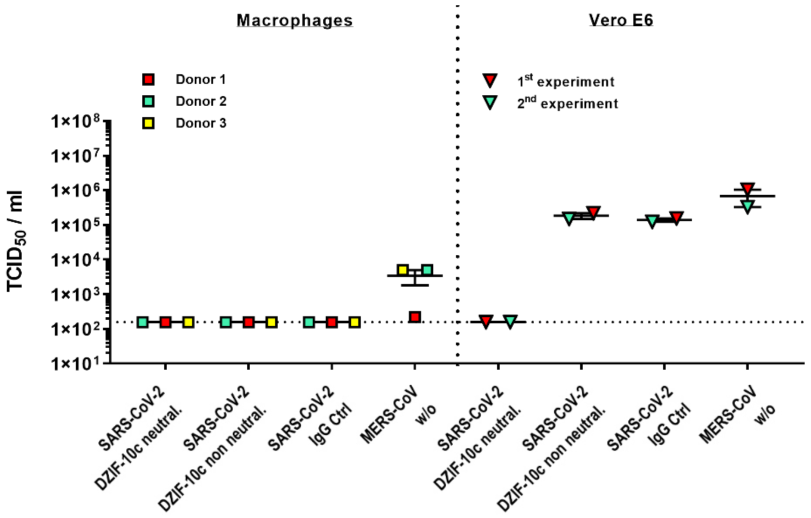


**Fig. S10:** ***In vitro*** **ADE Assay**

$CD14^+$-differentiated human blood macrophages were infected with SARS-CoV-2 in the presence of DZIF-10c at neutralizing and non-neutralizing concentrations or an IgG isotype antibody. General infectivity of macrophages was confirmed by infection with MERS-CoV. VeroE6 cells were included to confirm the adequacy of the selected antibody concentrations and viral infectivity. (A) SARS-CoV-2 and MERS-CoV genome copies in cell lysates determined by qRT-PCR. (B) Infectious SARS-CoV-2 and MERS-CoV titers in cell culture supernatants determined by TCID50 assay. Error bars represent mean ± SEM. Dotted lines indicate lower limit of detection.
